# Reproducible Propagation of Species-Rich Soil Microbiomes Suggests Robust Underlying Deterministic Principles of Community Formation

**DOI:** 10.1101/2021.10.29.466435

**Authors:** Senka Čaušević, Janko Tackmann, Vladimir Sentchilo, Christian von Mering, Jan Roelof van der Meer

## Abstract

Microbiomes are typically characterised by high species diversity but it is poorly understood how such system-level complexity can be generated and propagated. Here, we used soils as a relevant model to study microbiome development. Despite inherent stochastic variation in manipulating species-rich communities, both laboratory-mixed medium complexity (21 soil bacterial isolates in equal proportions) and high-diversity natural top-soil communities followed highly reproducible succession paths, maintaining distinct soil microbiome signatures. Development trajectories and compositional states were different for communities propagated in soils than in liquid suspension. Microbiome states were maintained over multiple renewed growth cycles but could be diverged by short-term pollutant exposure. The different but robust trajectories demonstrated that deterministic taxa-inherent characteristics underlie reproducible development and self-organized complexity of soil microbiomes within their environmental boundary conditions. Our findings also have direct implications for potential strategies to achieve controlled restoration of desertified land.

**TEASER:** Species-rich soil microbiomes grow and propagate reproducibly despite inherent stochastic complexity, paving the way for soil restoration.

## INTRODUCTION

Microbial communities are highly complex systems that self-organize seemingly spontaneously within the spatiotemporal, physical, chemical and biological boundary conditions of their environment or their host. The living microbial systems within these boundaries (the ‘microbiomes’) have attracted recent wide interest, due to their crucial contributions to ecological and biosphere processes (*1*–*3*), as well as to plant (*4*), human (*5*) and animal health (*6*). However, despite their widely recognized importance, there is still a large gap in understanding the general principles underlying microbiome development and functioning, as well as their amenability for functional and compositional engineering.

To a large part, our current understanding of the operating principles of microbiome formation comes from bottom-up studies with limited species numbers in synthetic ecosystems (*7*–*10*). Interspecific interactions are assumed to be the generators of community self-assembly and of emerging system-level metabolic properties (*11*, *12*). For example, range expansion experiments with 2-3 strains have demonstrated the quality, types and importance of interspecific metabolic interactions and spatial structuring (*13*–*18*). To some extent, higher-order community composition can also be successfully predicted from empirical measurements of paired growth interactions (*10*, *19*). However, multi-species interactions can give rise to feedback mechanisms that provide reciprocal control on their growth (*10*), or lead to multistable paths as a consequence of individual growth variation (*20*). Interspecific interactions further emerge in dependency of initial growth conditions and environments (*21*, *22*), and with increasing species complexity, non-additive effects may arise (*23*). The emergence of interspecific interactions depends on the spatial distance between cells (*24*) and, consequently, may be different in highly fractured environments such as soil, as opposed to liquid suspension (*25*–*28*). The question is thus whether developmental paths of species-rich communities are inherently stochastic and, in that sense, mostly irreproducible, or whether their taxa-composition provides robust self-organizing properties that will only diverge as a result of differences in environmental boundary conditions. In order to test this question, it is important to design studies that can bridge from the very simplified synthetic communities alluded to above, to more realistic species-diverse communities.

The major aims of the underlying work were thus twofold: first, to develop a tractable system to generate and propagate species-rich communities, and secondly, to study their developmental paths and resulting compositional states under different environmental boundary conditions and culturing regimes. We specifically focus on soil microbiomes, which comprise among the most diverse known microbial communities with up to 50 000 species (*29*) and 10^10^ cells per gram of material (*30*). The soil microbiome is of crucial importance for soil fertility and plant growth, for water purification and biogeochemical cycles (*1*, *31*, *32*). Soils are threatened world-wide as a result of land management, agricultural practices, erosion, waste deposition or chemical spills, leading to a general loss of soil structure and diversity (*33*, *34*). Soil microbiomes are thus highly relevant and one of the options for restoration of perturbed communities is through rational management, although current methods, e.g., soil transplantation or inoculation are very much a black box (*35*–*38*).

To obtain a realistic culturing system to propagate species-rich soil communities, we used autoclaved natural soil matrix replenished with soil carbon and nutrient extract. We contrast two types of soil communities, one composed of 21 indigenous soil isolates covering four major phyla (called *synthetic community* or SynCom), and the other comprising a species-rich soil microbial mixture directly washed and purified from top soil (NatCom, for natural community). Both communities were inoculated at low density in the reconstituted soils to allow growth and colonization, under two different culturing regimes. The first consisted of a single long-term incubated batch sampled after one week, two and six months, to favor slow-growing bacteria. The other consisted of multiple dilution-growth cycles of one week each, to favor community stabilization and test resilience to chemical perturbation. Community trajectories in soils were further compared to that in liquid suspension. Compositional changes were inferred from 16S rRNA gene amplicon sequencing and community signatures were compared to all available world-wide characterized soil and rhizosphere communities. Our results indicate highly reproducible species-rich community development for both synthetic and natural soil inocula. Developmental trajectories depend on incubation regimes and environmental conditions, suggesting robust deterministic self-organizing principles.

## RESULTS

### Design of controllable soil microbiome culturing systems

Standardized culturing systems for studying the development and succession of species-rich microbial communities were produced from sterile nutrient-complemented soils that could be inoculated, grown and diluted into fresh material, as is common for typical liquid culturing (Fig. 1A). The soil matrix was obtained from a riverbank sediment, twice autoclaved and complemented with a sterile soil nutrient extract from a topsoil (SE, soil extract). SE-reconstituted soils maintained general properties of natural soil. The final matrix of autoclaved soil + SE had a slightly basic pH, which was indifferent from the starting material (Fig. 1B). Autoclaving reduced Mg-levels, but did not significantly change Ca, P and Fe levels (Fig. 1C). UV/Vis fluorescence measurements indicated organics-rich material in SE (Fig. 1D, Supplementary table S1), which retained broadly the same six groups of humics and fulvics (*39*) as in the original soil (Fig. 1E, Supplementary table S2, p = 0.3078 Fisher’s exact test of relative abundances). Reconstituted soil showed increased availability of lower molecular weight compounds due to SE addition, as evident from the increased absorbance ratio at 250 nm to 365 nm (by 0.73 and 0.24 respectively; Fig. 1D, Supplementary table S1). Organic matter analysis in the SE-supplemented soils indicated an average of 1.5 mg total organic carbon g^−1^ soil matrix and 0.3 mg total N g^−1^ (Supplementary table S3). Assuming carbon needs of 200 fg C per cell and a g-C g-C^−1^ yield ratio of 20%, this would permit the development of a community of roughly 10^9^ cells g^−1^.

**Figure 1.**
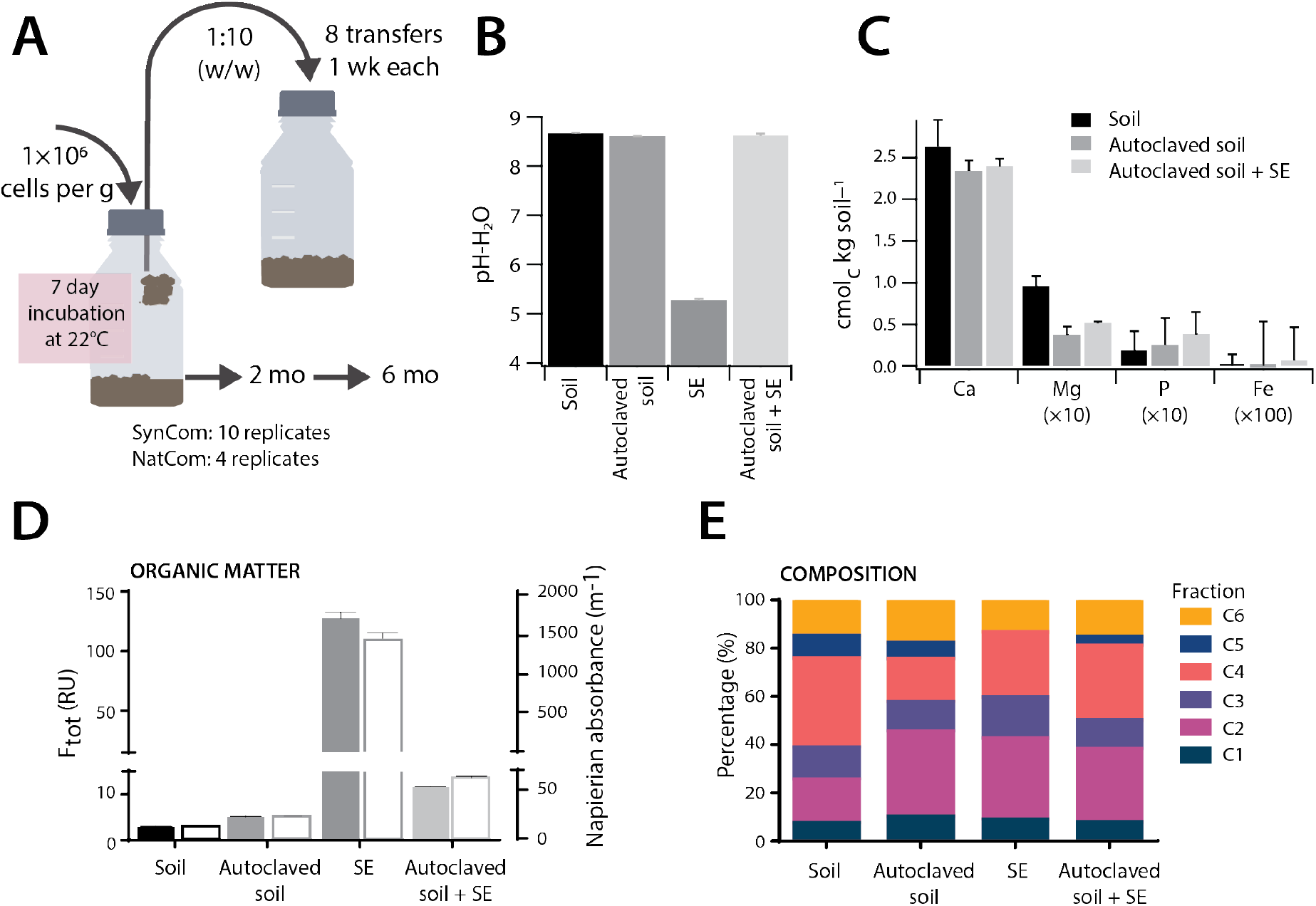
Characteristics of the soil culturing system. **(A)** Freshly washed soil communities (NatCom) or synthetic composed soil community (SynCom, 21 species) were used to inoculate 4 and 10 replicate sterile soil microcosms (each 100 g soil, 10^6^ cells g^−1^ at start), respectively. Microcosms were incubated for 7 days and then diluted into sets of fresh sterile microcosms (1:10, w/w). This growth cycling was repeated for a total of 8 cycles. The first microcosms were continued and sampled after 2 and 6 months. **(B)** Measured pH-H_2_O for soil before (black), after (dark grey) autoclaving and supplemented with SE (light grey), as well as the SE alone (grey). **(C)** Values of cation exchange capacity in cmol_c_ kg^−1^ of soil for soil before, after autoclaving and supplemented with SE. Bars show the mean from 3 replicates ± one *SD*. **(D)** Organic matter levels evaluated by UV/Vis measurements as total fluorescence (F_tot_) (filled bars, left axis, the same color scheme as in (C)) or Napierian absorbance (no fill, right axis, the same color scheme) for the materials used in the soil microcosms. Bars show means from 12 replicates ± one *SD*. **(E)** Inferred composition of organic matter by PARAFAC, according to the six defined groups of compounds in Ref. (*39*): C1, UVA Humic-like; C2, UVA-Humic like; C3-UVC-Humic-like; C4, Tyrosine-like; C5, UVA Humic-like and C6, Tryptophane-like. Bars are means from *n* = 12 replicate measurements.

### Generation and propagation of species-rich soil microbial communities

Autoclaved soil+SE was inoculated with starting community suspensions at 10^7^ cells ml^−1^, producing an equivalent of 10^6^ cells g^−1^ soil at the set 10% gravimetric water content. Community inocula consisted either of a washed and purified microbial cell suspension from top-soil (NatCom) or a suspension of 21 soil bacterial isolates mixed in equal relative abundances (SynCom, Table 1). Inoculated soil microcosms with NatCom suspensions after one week reached 2.8 ± 2.4 × 10^8^ cells g^−1^ (one *SD*, *n* = 4, Fig. 2A), an estimated 280-fold increase from the inoculum size (~8 doublings). Averaged across all 1–week culturing cycles, the NatComs maintained at 4.7 ± 1.1 × 10^8^ cells g^−1^ soil. This was an average of 3.5 times higher than the community size obtained in (liquid) SE alone (calculated on a per ml–basis, Fig. 2B, p = 0.0004, one-tailed t-test). This suggested that all easily accessible carbon was utilized during each week of incubation time and that communities reached semi-stationary phase (see below) before they were transferred to fresh soil medium. There was no discernable trend in the NatCom cell numbers as a function of growth cycle (Fig. 2A, NatCom linear regression: 0.0211, p=0.4371 compared to slope = 0). SynCom inoculum mixtures (Table 1) increased from 1×10^6^ cells g^−1^ soil to a stable average density after every growth cycle of 1.11 ± 0.32 ×10^9^ cells g^−1^ soil, which was 2.4 times higher than that of the NatCom (Fig. 2A, p = 9.9×10^−9^ unpaired two-sided t-test, *n* = 34). In comparison to its liquid SE suspension, the average one-week SynCom density in soil + SE was four times higher, similar as for the NatCom (Fig. 2B, p = 0.0002, one-tailed t-test).

**Table 1.** Taxonomy of selected strains for the synthetic soil community (SynCom).

| No. | Genus | Class | Phyla |
| --- | --- | --- | --- |
| 1 | **Microbacterium** | Actinobacteria | Actinobacteria |
| 2 | **Mucilaginibacter** | Bacteroidia | Bacteroidetes |
| 3 | **Curtobacterium** | Actinobacteria | Actinobacteria |
| 4 | **Variovorax** | Gammaproteobacteria | Proteobacteria |
| 5 | **Flavobacterium** | Bacteroidia | Bacteroidetes |
| 6 | **Cellulomonas** | Actinobacteria | Actinobacteria |
| 7 | **Tardiphaga** | Alphaproteobacteria | Proteobacteria |
| 8 | **Devosia** | Alphaproteobacteria | Proteobacteria |
| 9 | **Mesorhizobium** | Alphaproteobacteria | Proteobacteria |
| 10 | **Burkholderia** | Betaproteobacteria | Proteobacteria |
| 11 | **Pseudomonas** | Gammaproteobacteria | Proteobacteria |
| 12 | **Luteibacter** | Gammaproteobacteria | Proteobacteria |
| 13 | **Chitinophaga** | Bacteroidia | Bacteroidetes |
| 14 | **Lysobacter** | Gammaproteobacteria | Proteobacteria |
| 15 | **Pseudomonas** | Gammaproteobacteria | Proteobacteria |
| 16 | **Rhodococcus** | Actinobacteria | Actinobacteria |
| 17 | **Caulobacter** | Alphaproteobacteria | Proteobacteria |
| 18 | **Cohnella** | Bacilli | Firmicutes |
| 19 | **Rahnella** | Gammaproteobacteria | Proteobacteria |
| 20 | **Phenylobacterium** | Alphaproteobacteria | Proteobacteria |
| 21 | **Bradyrhizobium** | Alphaproteobacteria | Proteobacteria |

In contrast to the communities propagated under the 1–week growth/dilution cycles, those maintained under a single long incubation after initial growth in the first week, decreased in size as inferred from community DNA yields (Fig. 2C). NatCom DNA yields in soils decreased by 2–4 fold after 2 and 6 months, but not in liquid SE (Fig. 2C, p = 0.0101). SynCom sizes declined by 3– and 6–fold after 2 and 6 months, respectively, both in soils and liquid (Fig. 2C). This decrease may have been due to carbon limitation, consequent cell death and carbon turnover. Overall, these experiments indicated that high-density complex communities developed in both regimes and persisted over long times.

**Figure 2.**
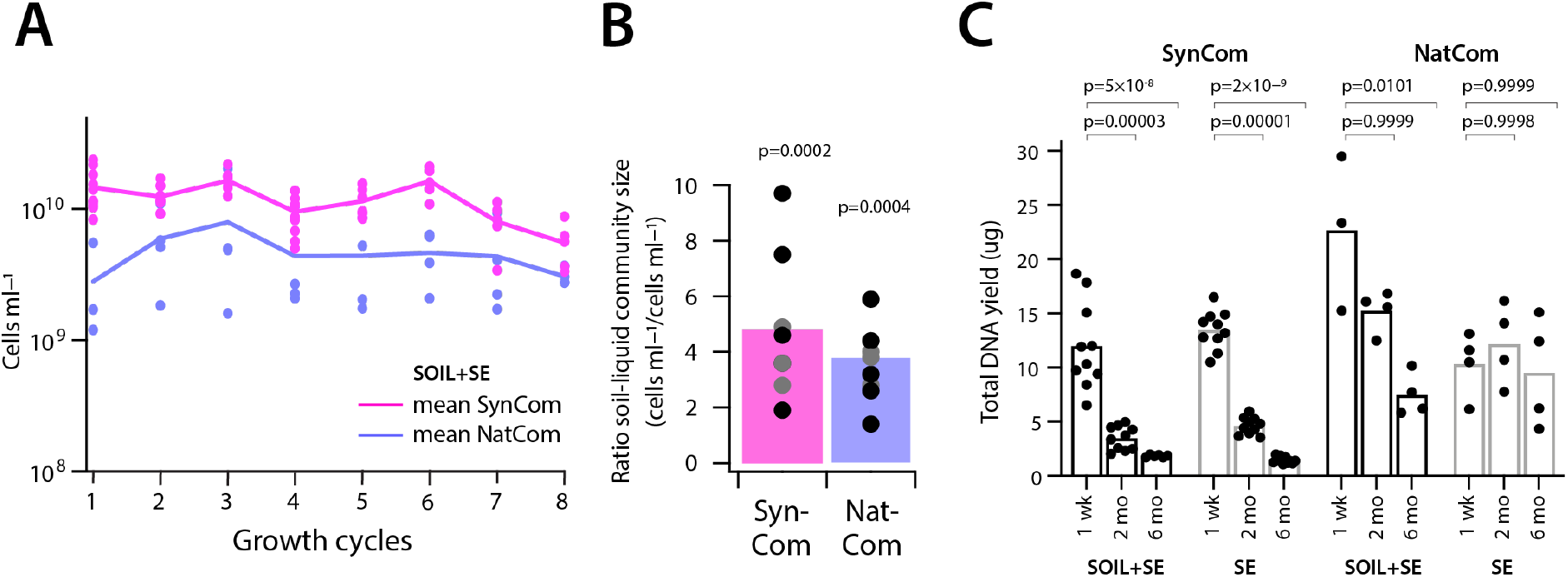
Development of synthetic and natural soil microbial communities in soil microcosms. **(A)** Sizes (in cells ml^−1^ soil liquid phase, determined by flow cytometry) of NatCom (cyan) and SynCom communities (magenta) across eight subsequent growth cycles. Lines connect the mean cell count of all replicates (NatCom: 4 replicates, SynCom: 10 replicates, except after the 5^th^ cycle where 5 replicates were removed for exposure to toluene) at the end of each transfer, with dots indicating individual values. **(B)** Mean (bars) and individual (dots, grey to black shades) for ratios of SynCom (magenta) and NatCom (cyan) flow cytometry cell counts after each growth cycle in soil + SE compared to suspended growth in liquid SE. P-values refer to one-tailed paired t-test of soil+SE values versus liquid SE suspensions. **(C)** Mean (bars) and individual (dots) replicate DNA yields from SynCom and NatCom communities after one week, 2 and 6 months incubation in soil + SE or in suspended growth in liquid SE. P-values refer to one-tailed paired t-tests in comparison to the 1–week DNA yields of the same sample group, with the *alternative* hypothesis that values at later time points are lower than week 1.

### Compositional state trajectories during culturing

The NatCom compositional dynamics under the two growth regimes was assessed from changes in the relative taxa abundances, determined by 16S rRNA gene amplicon sequencing using 99% identity thresholds for OTU assignment. The mean detected richness reduced from 233 in the inoculum to 22 (9%) after the first week, which slowly increased to 37 (16%) after the 8^th^ incubation cycle (Fig. 3A). In addition, 75% of the taxa after the 1^st^ cycle (week 1) were not detected in the inoculum (Fig. 3A, magenta bars), suggesting that community succession was initially driven by rapidly growing low abundant taxa. Non-metric multidimensional scaling (NMDS) analysis confirmed the strong deviation of both soil + SE and liquid SE microcosms from that of the original inoculum (Fig. 3B). Growth cycles resulted in closely clustering communities (Fig. 3B, T1–T8), whereas the single long incubations showed succession and higher similarity to the inoculum state (Fig. 3B, 2 and 6 months). NatCom development in soil + SE was distinct from that in liquid SE alone, indicating that the soil environment (and resulting pH differences) may have driven the community differentiation (Fig. 3B, adonis p = 0.001 with beta-dispersion of p = 4.38 × 10^−10^).

**Figure 3.**
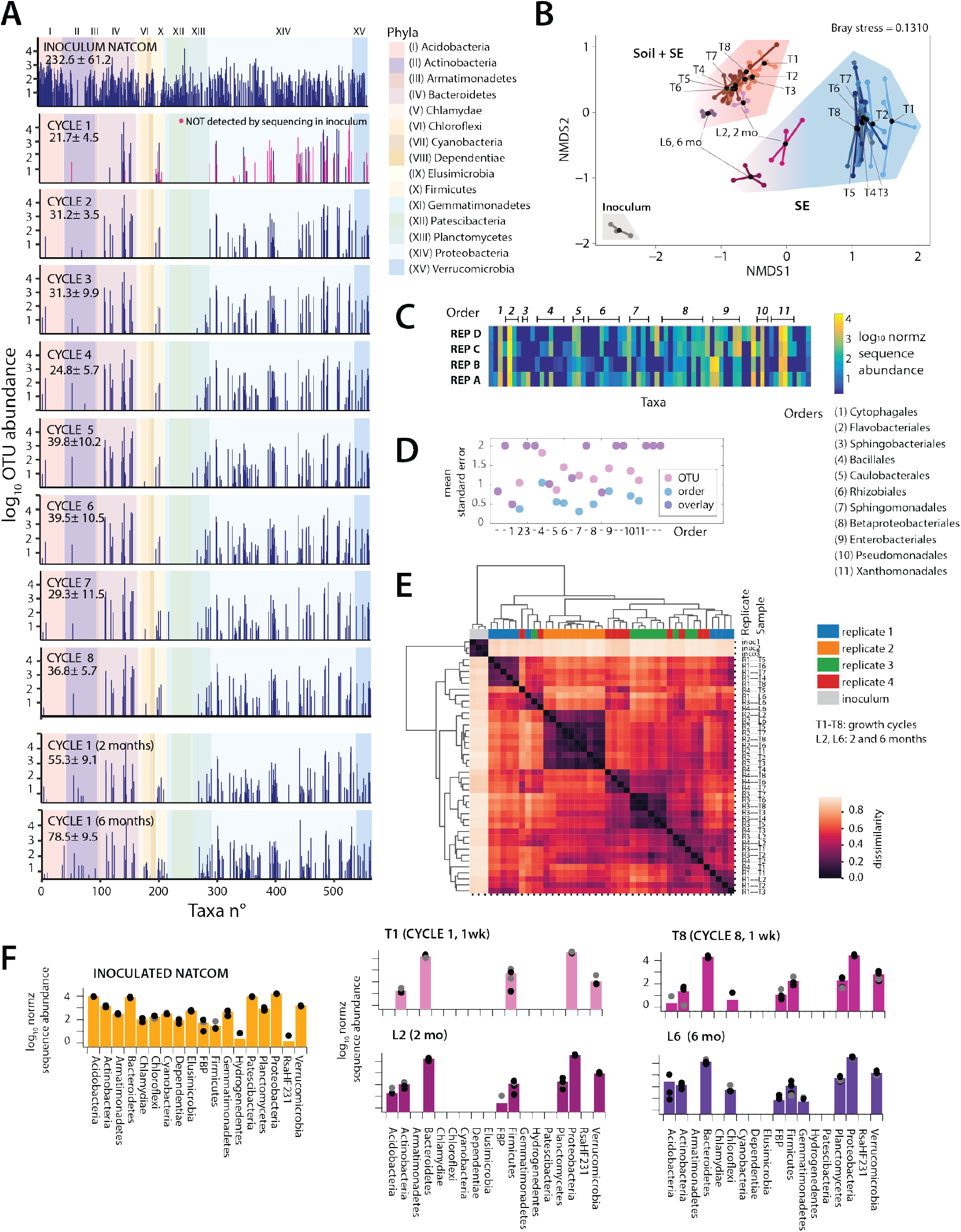
Community succession and composition of NatComs. **(A)** Mean log_10_-transformed total read-normalized (5 × 10^4^) taxa abundances in the soil inoculum, after all the 8 one-week growth cycles, and in the longer-term incubation timepoints (2 and 6 months). Abundance bars positioned according to taxa numbering from the OTU list (SILVA, above 99% similarity), with background color representing phyla affiliation (Roman numbering, according to legend). Numbers within panels show mean taxa richness ± one *SD* (*n* = 4 replicates). Magenta bars in the CYCLE-1 data point to taxa not detected in the inoculum. **(B)** Non-metric multidimensional scaling ordination of NatCom succession in soil + SE (magenta area), or in liquid SE suspension (cyan area; T1–T8, weekly transfers; L2, L6, two and six months incubations). Ordination plot based on Bray-Curtis distances. **(C)** Compositional variation (shown as log10-normalized abundance heatmap) among the four NatCom replicates (REP A–D) after the first growth cycle. Numbers above refer to taxa within order-levels as specified on the right. **(D)** Mean standard error of replicate variation (REP A–D after one week) at OTU-level (mean of means grouped within corresponding order, pink) or at order-level (blue; purple is where both OTU- and order-values overlap). Note how order-level variation is lower than OTU-variation. Numbers refer to order in (C). −, single OTU in order; not specified. (**E**) NatCom pairwise sample comparison, clustered by average-linked Bray-Curtis distances (color scale). Inoculum (soil 1-3) and replicates (R1–4) are highlighted by different colors on the top, growth cycles (T1–T8) or long-term incubations (L2, 2 mo; L6, 6 mo) in small fonts on the right. Note the strongly maintained replicate signatures (e.g., replicate 2). **(F)** Mean (bars) and individual replicate (grey to black dots, *n* = 4) grouped phyla composition of NatCom inoculum (orange), after the first growth cycle (CYCLE 1, 1 wk), the 8^th^ (CYCLE 8), and after 2 and 6 months (mo).

Although replicates clustered coherently in NMDS (Fig. 3B), there were obvious stochastic effects of compositional succession, illustrated by variation in appearance and relative abundance of individual taxa among replicate inoculations after the first week of incubation (Fig. 3C, e.g., Rhizobiales, Sphingomonadales and Enterobacteriales). Replicate variability was higher at OTU level than at order level (Fig. 3D, Supplementary figure S1), suggesting conserved functional order traits that permit strains from such groups to quickly colonize new environmental niches. NatCom replicates kept a relatively strong individual signature independent of multiple growth/dilution cycles (most evident with the “–2” replicate, Fig. 3E), which mostly converged in long-term incubations (Fig. 3E, L6 samples). This might be due to initial stochastic compositional variations that influence growth in the first incubation and from there on, propagate the states of regrown communities. Mathematical simulations of community growth and composition suggested that this variation may be due to subsampling effects of rare taxa with high growth rates within a finite-sized inoculum (Supplementary figure S2). Initially composed of 18 phyla, only five were detected in NatCom replicates after the first growth cycle, and four more appeared after cycle 8 (Fig. 3F), indicating that their members were present but undetectable at our sequencing depth. In contrast, long-term incubated NatCom showed members of ten phyla, indicating that this growth regime permitted higher diversity, perhaps by avoiding bottlenecks of the dilution/growth cycles on slow-growing members (Fig. 3F). This showed that species-rich soil communities can be grown and maintained with relatively constant composition over multiple dilution cycles, despite having inter-replicate stochastic strain variability. Culturing in SE-reconstituted soil clearly provided additional benefits to the community, since both its size (Fig. 2B) and its richness remained larger (by 12.02% with growth cycles and 9.31% in the long batch regime, Supplementary figure S3) than that in SE liquid suspension.

### Development of medium complexity synthetic soil microbiome recapitulates natural states

To corroborate the observed succession and development patterns in NatCom, we followed changes in the defined SynCom. The SynCom was selected from a total of 172 recovered pure cultures based on different colony morphologies and growth characteristics grouped into 52 different genera belonging to four phyla (Supplementary table S4). Except for the phylum Verrucomicrobia, these isolates covered the major phyla observed after the first NatCom growth cycle (Actinobacteria, Bacteroidetes, Firmicutes, and Proteobacteria, Fig. 3F), with some redundancies (Table 1, Fig. 4A).

**Figure 4.**
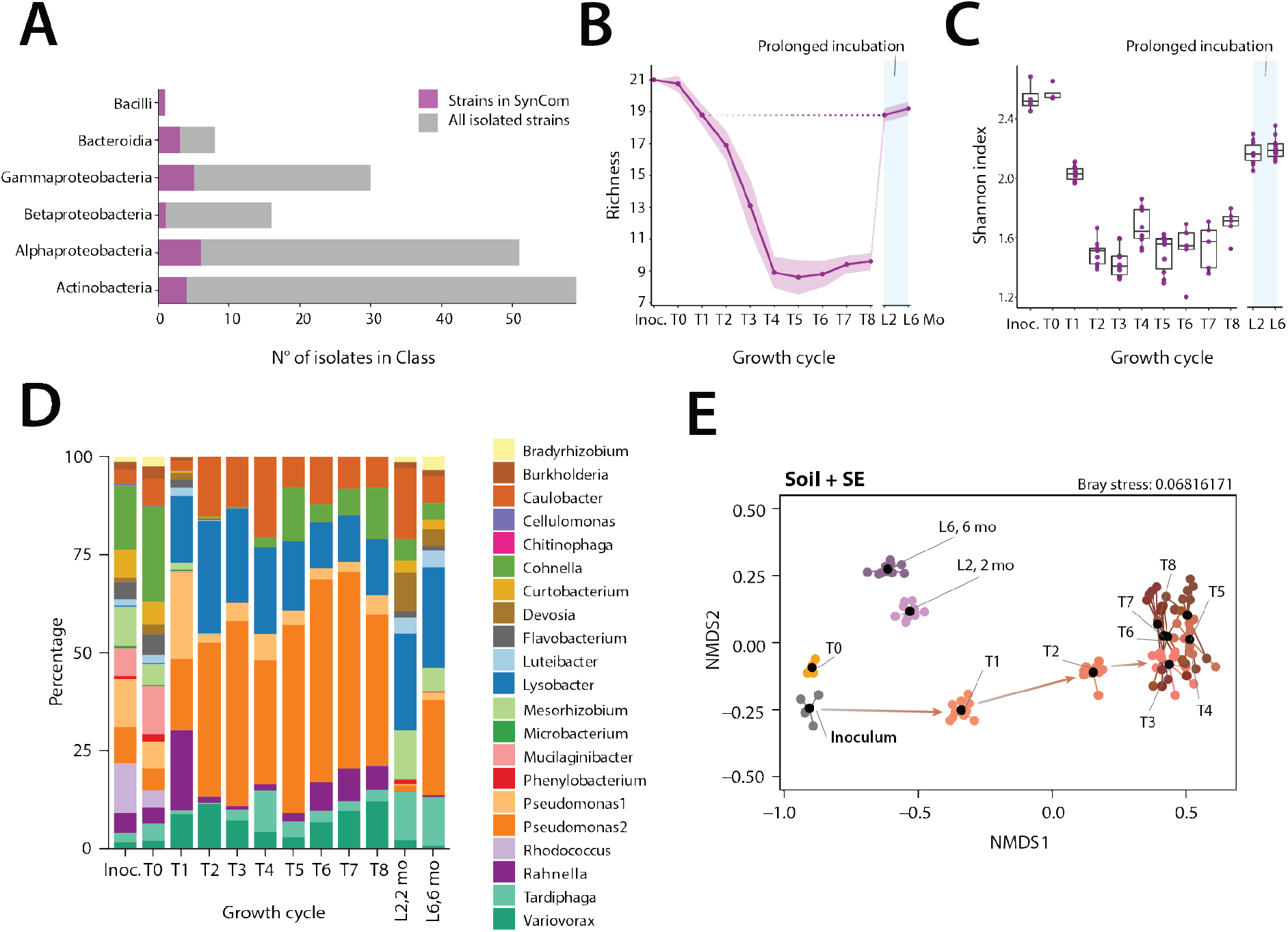
Succession and stabilization of a synthetic soil community over multiple growth cycles and long-term incubation. **(A)** Class attribution of the 172 isolated soil bacterial strains, and of the selected 21 strains of the SynCom. **(B)** Changes in mean SynCom richness (magenta line, ± one *SD* in light color background, *n* = 10 replicates) and (**C**) in mean Shannon indices (box plots, *n* = 10; except T6-T8; *n* = 5 replicates) throughout the 8 growth cycles in soil + SE (T1–T8), and during long-term incubation (L2, L6; 2 and 6 months, mo). **(D)** Stacked mean relative abundances (in percentage, *n* = 10 replicates) of SynCom members (legend on the right) from inoculation to the last growth cycle, and upon long-term incubation. **(E)** Non-metric multidimensional scaling of normalized SynCom compositions according to their Bray-Curtis distances. Black dots show the community centroids; colored dots are individual replicates.

In contrast to NatCom, the compositions of the SynCom showed clearer succession during the first three growth/dilution cycles, after which they stabilized. This was evident from a loss of apparent diversity (i.e., within the sequencing threshold for community membership), from 21 to 9–10 detectable members after the fourth cycle (Fig. 4B), and a sharp decrease of Shannon index (Fig. 4C). The T_0_ –sample (taken 30 min after inoculation into the soil) resembled the inoculum closely (Bray-Curtis distances of 0.26 ± 0.02, while the distance between inoculum and T8 was 0.65 ± 0.03), showing minimal bias introduced by cell extraction (Fig. 4D). Initially higher relative abundances of *Pseudomonas* strain1 and *Rahnella* during the first-to-third growth cycles were replaced by *Pseudomonas* strain 2, *Lysobacter*, *Variovorax*, and *Caulobacter* as the dominant members. Finally, also *Cohnella*, *Rahnella* and *Tardiphaga* regained sizeable proportions of the SynCom (Fig. 4D). Independent SynCom replicates followed highly similar developmental paths (Fig. 4E), in terms compositional changes, loss of diversity and reaching semi-stable compositions after the 4^th^ cycle (Fig. 4B-D). SynCom replicates clustered coherently over time and did not maintain individual replicate signatures as NatCom (Supplementary figure S4). SynCom compositions in soil + SE differed significantly from that of the inoculum and those grown in liquid SE suspension (Supplementary figure S5; adonis p = 0.001; betadisper p = 0.0002). Similar as for the NatCom, the long incubation regime led to higher detectable diversity of 18-20 strains from the initial 21 after 2 and 6 months (Fig. 4B, C, and E, p = 0.001 from adonis and p = 0.0002 for beta-dispersion). This included higher relative abundances of *Mesorhizobium, Luteibacter* and *Devosia* compared to e.g., *Pseudomonas* (Supplementary figure S5).

### SynCom and NatCom retain soil signatures but differ in replicate variability

In comparison to a wide set of available soil communities (*n* = 110,928), both SynCom and NatCom compositions grown in soil kept clear soil community signatures (Fig. 5A). Interestingly, SynCom compositions located closer to ‘plant rhizosphere’ communities, possibly due to the culturing isolation bias (Fig. 5A). NatCom grouped closer to ‘field soils’, whereas the inoculum, as expected, had a ‘forest’ soil signature (Fig. 5A). There is not a clear single factor underlying this environmental signature, although soil-pH (as far as present in the meta-data) seems an important variable (Fig. 5B). Both SynCom and NatCom became largely dominated by Alpha- and Gammaproteobacteria, but were notably different in the relative abundances of Bacteroidetes (contributing 30-50% in the NatCom) and Firmicutes (5-10% in the SynCom) (Fig. 5C). SynCom replicate variability was twice as low as that of the NatCom (Fig. 6D, F=17.495, p=5.19 ×10^−5^, ANOVA), with high replicate homogeneity (i.e., the replicate Bray-Curtis distance from the community centroids, ranging from 0.01 to 0.20; Fig. 5D). The reason for this is likely the lower number of starting strains in the SynCom and lower likelihood of stochastic variations as a result of subsampling upon dilution (as in, *e.g*., Supplementary figure S2).

**Figure 5.**
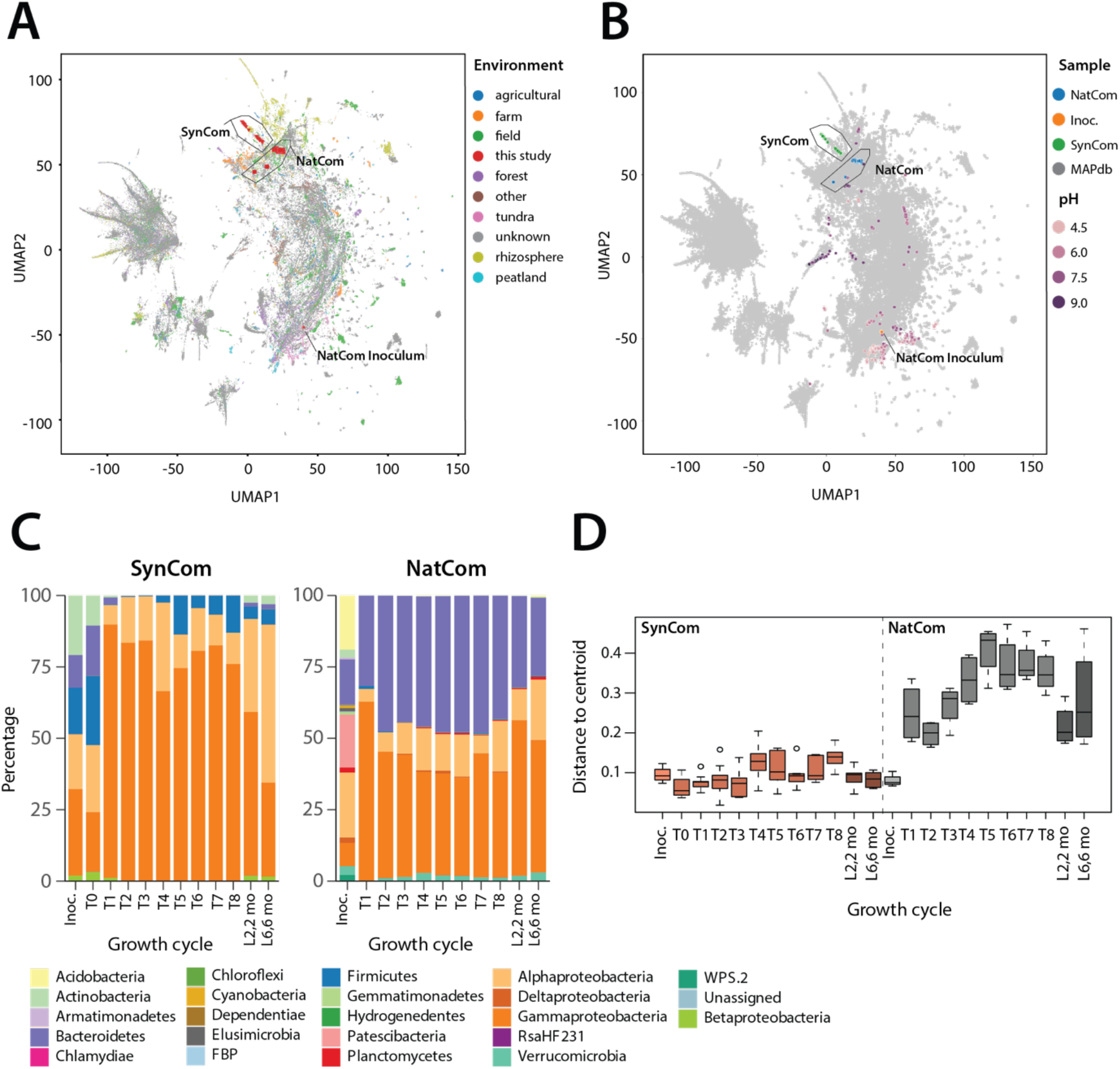
NatCom and SynCom community signatures. **(A)** Environmental signature of NatComs and SynComs. Map shows a UMAP projection of SynCom and NatCom samples together with 110,928 soil communities (dots) extracted from the Microbe Atlas Project (*70*), based on Bray-Curtis distances and color coded along their environmental origin, or **(B)** overlaid with soil pH, extracted from the Earth Microbiome Project (*71*). **(C)** SynCom and NatCom relative abundances at phyla and class levels (Proteobacteria only). **(D)** Interreplicate variability of SynCom and NatCom replicates, shown here as individual Bray-Curtis distances to the corresponding community centroid. Boxplots show 25^th^, median and 75^th^ percentiles, with whiskers indicating 1.5× the interquartile range.

**Figure 6.**
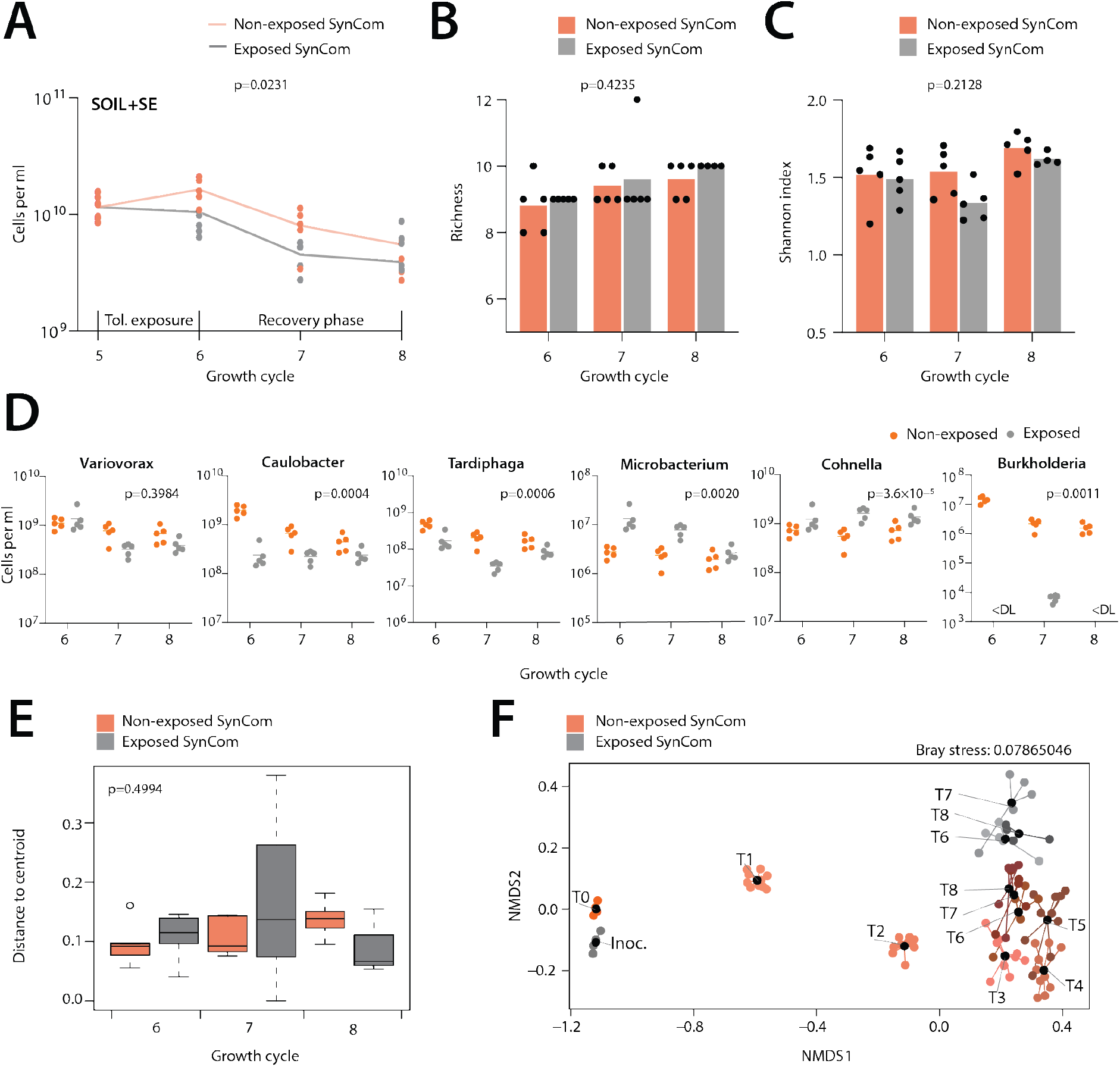
Community resilience upon chemical perturbance. **(A)** Mean (lines) and replicate (dots) SynCom size changes (in cells ml^−1^ soil liquid phase) in toluene-exposed (grey) versus non-exposed (orange) microcosms (each 5 replicates). Toluene exposure during 1 week of the fifth growth cycle. P-value refers to comparison of cell densities after the 6^th^-8^th^ cycles between exposed and non-exposed communities in a Wilcoxon matched-pairs signed rank test. **(B)** Mean (bars) and replicate (dots) richness in toluene-exposed versus non-exposed SynComs. P-values from two-way ANOVA for toluene exposure. **(C)** as B but for Shannon index. **(D)** Changes in absolute abundances (calculated from individual relative sequence abundances and total community size by flow cytometry) of selected SynCom members with and without toluene exposure (two-sided t-test, grouped T6–T8 values). **(E)** Interreplicate variability, expressed as average distance of replicates to community centroid. P-value from ANOVA of exposed versus non-exposed centroid distances. **(F)** Toluene-exposure effect on SynCom compositions after the 6^th^-8^th^ growth cycles (NMDS based on Bray-Curtis community distances). Sample abbreviations (e.g., T1) as before; black dots, community means; grey dots, toluene-exposed SynCom replicates during T5; orange to brown dots; non-exposed SynCom.

### Chemical perturbation changes SynCom trajectories

In order to investigate the stability of developed communities, we tested their resilience towards the moderate toxic compound toluene, as an example of recurrent soil pollution with organic solvents (*40*). To this end, we split the stabilized ten SynCom replicates in two groups of five after the fifth growth cycle; one series of which was exposed to toluene vapor during the next one-week cycle, the other cultured as before. After this exposure period, all SynCom replicates were diluted again into sterile, non-polluted SE-amended soil; and growth cycles were continued as before. Exposure to toluene significantly lowered the attained community sizes (Fig. 6A, p = 0.0231, one-tailed t-test on all replicate samples and time points, n = 15). In contrast, toluene exposure did not lead to significant changes in richness (Fig. 6B, p = 0.4235, two-way ANOVA), nor did it influence Shannon diversity (Fig. 6C, p = 0.2128, two-way ANOVA). Varying effects were observed on individual SynCom members, which either slightly (Fig. 6D, e.g., *Variovorax*, 25.8% decrease), or drastically decreased in population size (Fig. 6D, 99.9% decrease of *Burkholderia*, see *Devosia* and *Flavobacterium* in Supplementary figure S6), whereas some increased in abundance (Fig. 6D, *Microbacterium*, *Cohnella*). Inter-replicate variability was not significantly affected with toluene exposure, even during the first week of recovery (Fig. 6E, F=0.8973, p=0.4994, ANOVA). Community signatures in exposed SynCom remained distinct from those of the non-exposed communities even after the 8^th^ cycle (Fig. 6F, adonis p = 0.001; betadisper p=0.1024). Altogether, this indicated that stable soil communities can be perturbed by chemical (toluene) exposure, which changes their compositional trajectories in a long-lasting manner.

## DISCUSSION

We showed reproducible assembly, succession and composition of both a high-complexity NatCom (starting from washed mixed soil inoculum, containing 18 bacterial phyla), as well as a medium-complexity SynCom with 21 culturable strains (covering four major bacteria phyla) in a soil culturing system that enables soil-to-soil transfers. The standardized soils were generated from riverbank sediment supplemented with a soil extract, which provided realistic available carbon and nitrogen compounds for growth of community cell densities similar to what is typically found in top soils (*30*). Both NatCom and SynCom retained typical soil and plant rhizosphere microbiome signatures, and thereby represent excellent test beds for plant growth, community management or soil resilience studies that require complex and reproducible starting communities. Both growth regimes, either imposed as multiple one-week growth and dilution cycles in soil, or as single batch long incubation (up to 6 months) favored establishment of high species diversity, which in short incubations (one week) was dominated by relatively fast-growing opportunistic strains. Cultured SynCom on average had higher cell densities on the same substrate than NatCom, which might be due to their inoculum being exclusively of bacterial origin, without any potential phage or protist predators that might have been present in the washed mixed microbial top-soil NatCom inoculum mixture. The sizes of both communities, however, moderately decreased over long incubation periods (2 and 6 months), suggesting some cell death and consequent carbon turnover. The long-time incubation may have allowed growth of members from slow-growing phyla that are difficult to obtain in pure culture (e.g., Acidobacteria, Gemmatimonadetes).

One of the key surprises of our work is the demonstration of highly reproducible trajectories and compositional states of medium-to-high species-diverse soil communities. Low variability among starting replicate soil communities makes it easier to detect effects of inoculant bacteria or fungi in relation with plants for improving plant health (*41*), to investigate the influence of bioaugmentation agents in pollution removal (*35*, *36*), or to address fundamental ecological questions on community resilience (*42*). These growth properties of SynCom and NatCom can be fruitfully exploited in future work to study fundamental ecological questions of species redundancy, resilience, or invasion resistance (*42*, *43*). The high reproducibility of community development under realistic culturing (e.g., soil) was counterintuitive. Considering the complexity of the provided nutrients, highly fractured soil environment and species-diverse inocula, we expected that stochastic small variations in experimental manipulations would lead to chaotic system behaviour. Contrary to this intuition, the medium complexity (21-member) SynCom developed highly reproducibly among ten replicates, with similar succession patterns, total community sizes and relative species abundances. Initially (inoculated) balanced species proportions were quickly replaced by coherent compositional trajectories and states during multiple growth/dilution cycles. The attained SynCom compositional states were dependent on the type of incubation regimes (growth cycles versus one-batch long time), and specific environments (soil+SE or liquid SE) but retained no invidual replicate signatures. Compositional states could be perturbed by short-term chemical exposure, but then reproducibly continued on slightly different trajectories. In contrast, NatCom compositions showed more stochastic variability among replicates, and individual replicate signatures were retained to some extent in the growth/dilution cycles. Community growth simulations suggested that the reason for increased stochastic replicate variability may lie in population bottlenecks arising from finite-sampling of high diverse community inocula with rapidly growing colonizers, leading to a state which then self-propagates in subsequent growth/dilution cycles. Despite this, NatCom replicate variability collapsed at a higher phylogenetic level, suggesting similar functional and redundant properties in the complex starting inoculum that are selected during colonization of pristine growth environments. Long-time incubations also dissipated NatCom compositional variations to a large extent. Both SynCom and NatCom in soil microcosms developed and maintained clear soil community signatures. This indicates strong deterministic influences of the initial species composition on the community development trajectories (i.e., self-organizing complexity) within its system boundaries and the prevailing environmental conditions.

Several authors have reiterated that the origins of microbiome complexity remain fundamentally unknown (*23*, *44*) and that general rules governing community assembly and functioning are difficult to deduce (*21*, *45*–*47*). It is clear that community growth and development are influenced by a myriad of factors such as growth substrates, spatial structures, and presence of other chemical compounds (*44*, *48*). The more complex carbon substrates deployed in this study possibly require and facilitate a wider range of metabolic capacities and therefore maintained higher functional diversity (30–40 OTUs in NatComs), than in previous experiments starting with soil and phyllosphere communities but grown on a single carbon substrate (5–12 exact sequence variants) (*49*). Community development is further expected to be dependent on emerging interspecific interactions leading to transcending systems-level functionalities (*21*, *45*). Indeed, both NatCom and SynCom development seemed strongly determined by their starting taxa compositions, on top of which the environmental boundary conditions (i.e., soil versus liquid) influencing the community trajectories. The difference in compositional trajectories and states in soil and liquid, despite containing the same complex nutrient resource availability (soil extract), may be due to different types or magnitudes of interspecific interactions arising in the spatially structured, disconnected and heterogenous growth environment of the soil as opposed to the liquid-suspended growth (*25*). Soils are expected to provide unique ecological niches (*1*, *50*), and their aggregates affect nutrient availability and gradients in electron donors and acceptors (*26*–*28*, *48*, *51*). Indeed, SynComs and NatComs maintained on average higher species diversity in soil microcosms than equivalent liquid cultures, suggesting emerging favorable dependencies, which permitted more phyla to sustain and grow (*25*). Suggestive for this is that members belonging to the Acidobacteria, Verrucomicrobia and Planctomycetes proliferated in all NatCom microcosms, whereas we did not manage to culture them individually using the same nutrient substrates.

Natural soil communities probably only very rarely have the opportunity to colonize a pristine soil environment, except perhaps for soil transplants or soil construction work, grubbing, glacier retreats or other (*52*, *53*). At a large scale (cm – m), the composition of complex natural soil communities is stable, but may undergo temporal and very local fluctuations driven by nutrient gradients from plant roots, burrowing fauna, rainfall, seasonal temperature changes or other (*54*–*56*). In that sense, our long-term incubation regimes resembled new soil colonization events, eventually leading to a mature state composition, typically comprised of several abundant members and a vast fraction of extremely low abundant species (“rare biosphere”) (*57*–*59*). The regime of imposed growth cycles may reflect what happens at sudden bursts of newly available carbon in the soil. As the NatCom experiments demonstrated, some “rare taxa” in the mature compositional state as isolated from the natural soil (Gammaproteobacteria, known generalists) rapidly proliferated in the first week of incubation, with Alphaproteobacteria and other phyla appearing only later, as has been observed before in natural systems (*47*, *59*). Some rare taxa may thus rather represent “conditionally rare taxa”; those with radically changing abundances depending on space and nutrient availability (*60*). The specific roles or capacities of those taxa to become more abundant over time remain unclear for now, and could be due to factors such as use of different (refractory) carbon substrates, predatory lifestyles, different nutrient requirements or forms of metabolic dormancy to remain viable for longer. From an engineering perspective the maintenance of temporary community compositional steady states by the cycling growth-dilution regime is interesting and suggests an avenue for approaches that aim to keep relatively constant species proportions in mixed communities over time. Reproducible propagation of soil communities will also be key for restoration efforts on degraded or desertified land that aim to bring back healthy soil life.

## MATERIALS AND METHODS

### Preparation of a natural soil community

A natural mixed microbial community (NatCom) was washed from batches of 20 g taken with a sterile metal spoon from the 5 cm topsoil layer after removal of twigs, roots and leaves (Dorigny forest, University of Lausanne, 46°31’16.4”N 6°34’43.0”E). Soil batches were immediately transported to the lab and processed within 1 h. The soil was sieved through a 3–mm mesh to remove large particles. Microbial cells were detached from soil particles by mixing with sterile 0.2% (*w*/v) tetrasodium-pyrophosphate decahydrate solution (pH 7.5, Sigma-Aldrich), and then purified by sucrose gradient solution centrifugation as described by (*61*). The cell suspension recovered after sucrose gradient centrifugation was twice washed with sterile saline solution (0.9% NaCl) and resuspended in the same. Serial dilutions were stained with SYBR Green I and cell numbers were counted using flow cytometry (see below). For inoculation into microcosms, the cell suspension was diluted in soil extract (SE, see below) to 10^7^ cells ml^−1^. Subsamples of the NatCom suspension were used for DNA extraction and 16S rRNA gene amplicon sequencing (see below).

### Preparation of the synthetic soil community

Individual soil isolates were obtained from similar NatCom suspensions of the same soil location, additionally purified using Nycodenz gradient (*61*), diluted and plated on different media, as suggested by Balkwill and Ghiorse (*62*). We used PTYG medium (containing, per L: 0.5 g glucose, 0.5 g yeast extract, 0.25 g peptone, 0.25 g trypticase, 0.6 g MgSO_4_·7H_2_O, 0.07 g CaCl_2_·2H_2_O, 15 g agar), or soil extract medium (see below) solidified with 1.5% agar (Agar bacteriological, Difco), either at pH 4.5 (adjusted with hydrochloric acid) or at pH 7.5 (with sodium hydroxide). All plates were incubated at room temperature (23 °C) for 2 weeks. In total, 172 morphologically distinguishable colonies were selected, purified to homogeneity by streaking on the same medium, regrown in PTYG and stored in 15% (*v*/*v*) glycerol at –80°C. Strains were identified and taxonomically positioned by full length 16S rRNA gene sequencing (see below).

A set of 21 isolates representing different major phylogenetic and culturable groups (Table 1) were selected to assemble a synthetic soil community (SynCom). To prepare the SynCom inoculum, individual strains were plated from −80° stocks on PTYG agar and grown for 4 days at room temperature. Cells were then collected from the plates by washing with 5 ml of soil buffer (containing per L, 0.6 g of MgSO_4_·7H_2_O, 0.1 g of CaCl_2_ and 1.8 ml of 5 x M9 minimal salts solution [BD Biosciences]). Individual cell suspensions were serially diluted in soil buffer and stained with SYBR Green I for 15 min in the dark, according to manufacturer’s instructions (Invitrogen), after which cell numbers were counted by flow cytometry (see below). Pure cultures were then diluted in soil extract (SE, see below) and mixed to obtain a suspension of in total 10^7^ cells per ml, and with approximate equal abundances of each individual member.

### Soil microcosm preparation

Both NatComs and SynComs were cultured and passaged in semi-natural sterile soil systems, based on a coarse silt supplemented with a sterile soil extract solution. The soil matrix was prepared from riverbank sediment (0-10 cm horizon) of the Sorge river sampled at the campus of the University of Lausanne (46°31’22.4”N 6°34’31.7”E). The material was transported to the laboratory, spread in 5 cm layer in trays and air-dried in a ventilated hood at 23°C for two weeks, followed by double sieving to retrieve the 0.5-3 mm sized soil fraction. Sieved soil was divided in 2 kg portions, autoclaved for 1 h at 120°C and dried for an additional seven days as described above. Batches of soil (90 g for the first inoculation series, 80 g for subsequent transfers) were then distributed into 500–ml Schott borosilicate glass flasks with plastic screw cap and seal. Individual flasks with soil were again autoclaved (20 min, 120°C) to kill any remaining spores and vegetative cells. The sterility of the soil was confirmed after the second autoclaving by sampling batches of 10 g with a sterile glass spoon, mixing with 20 ml sterile 0.2% pyrophosphate solution and vortexing for 1 min at maximum speed (Vortex-Genie 2, Scientific Industries, Inc.), after which aliquots of 100 μL were plated on three different agar media: PTYG (see above), R2A (DSMZ GmbH) and Nutrient Agar (BD Biosciences). Absence of grown colonies after 3 weeks incubation at room temperature was taken as indication for the material to be sterile. All microcosms used in the study originated from the same batch of sieved soil.

As source of nutrients for all microcosms we produced a soil extract (SE) from the same soil as used for the NatCom and the SynCom isolates (see above), as follows. Top soil material (1–5 cm layer, 6 kg) was sampled as before and mixed in a 1:1 volumetric ratio with tap water in batches of 2 kg. The mixture was autoclaved (1 h, 120°C) mixed and left to settle overnight. The resulting supernatant was decanted into sterile 250 ml centrifuge tubes, centrifuged at 5000 × *g* for 15 min to remove solids and pooled into 500 ml Schott flasks. This solution was autoclaved once more and then filtered through a 0.2−μm Stericup Quick Release System PES filter (Merck) into clean sterile Schott glass flasks and stored at room temperature in dark. The pH of SE was 5.28 ± 0.03. Its total organic carbon content (TOC) equaled 753 ± 49 mg C l^−1^. A single batch of SE was used for all microcosms in this study. Analysis of soil parameters is described in the Supplementary methods.

### Soil microcosm inoculation and culturing

Soil microcosms were inoculated with NatCom (four replicates) and SynCom (10 replicates) suspensions, and cultured either as a long-term single batch incubation, or through multiple one-week growth and dilution cycles (Fig. 1A). Each microcosm initially comprised 90 g dry sterile soil matrix in a 500 ml screwcap glass bottle, amended with 10 ml community inoculum (at 10^7^ cells ml^−1^ in SE, see above), thus resulting in ca. 10% gravimetric water content and 10^6^ cells g^−1^ soil at start. The pH(H2O) of the soil microcosms after inoculation was 8.62 ± 0.04. Uninoculated soils (four replicates) amended with 10 ml sterile SE served as controls for potential contamination. To contrast community growth in liquid suspension, the same SynCom and NatCom inocula were grown directly in 10 ml SE in 50 ml sterile Falcon tubes (starting at 10^7^ cells ml^−1^), which were incubated at ambient temperature in the dark. After inoculation and before each sampling the soil microcosms were thoroughly mixed on a horizontal roller mixer (20 min at 80 rpm). SE-liquid microcosms were vortexed for 1 min every day.

In the long-term incubation series, samples (20 g) for community analysis (see below) were taken from each replicate microcosm after one week, two and six months. In the cycling regime, 11 g of the microcosm material were aseptically transferred after one week of growth to a fresh flask containing 80 g of dry sterile soil matrix. 9 ml of sterile SE was again added to maintain moisture content and replenish nutrient levels, thus resulting in ten-fold microcosm dilution upon each transfer. Flasks were again incubated for 1 week as before with intermittent roller-mixing. This incubation-dilution cycle was repeated eight times consecutively.

SE-liquid microcosms were sampled (2 ml) each week for community analysis, after which 1 ml was transferred to a fresh tube with 9 ml of sterile SE. Incubation and dilution were repeated for eight cycles, similar as for the soil microcosms with the cycling regime. A further SE-liquid control was prepared for the long incubation (one week, 2 and 6 months).

### Chemical perturbation

In order to assess the effect of chemical perturbance on the resilience of the established communities, five of ten SynCom replicates (both soil+SE and SE-liquid) after the fifth transfer (see above) were exposed to toluene vapor during one week, as follows. After the inoculation with material from the previous cycle, heat-sealed 1 ml (for soils) or 0.2 ml (for SE-liquid) micropipette tips were placed inside the microcosms, open at the top to the air, and filled with 100 μl or 10 μl pure toluene, respectively. These volumes are equivalent to a nominal concentration of 1.88 mM toluene, which will partition into the gas and aqueous phases in both systems. Microcosm flasks and tubes were tightly closed and incubated for 7 days with daily mixing (during each mixing, the toluene reservoir was briefly removed and then placed back). Samples were taken at day 7, and material from the exposed microcosms was again diluted as before into fresh soil+SE or SE-liquid, but without toluene. The non-exposed growth regime was repeated for another two cycles to study community recovery.

### Community analysis

Samples of 20 g (soil+SE) or 2 ml (SE-liquid) were mixed with 20 ml of sterile pyrophosphate solution (see above) and vortexed for 1 min at maximum speed. The samples were left to stand for 1 min to settle soil particles, after which the supernatant was transferred aseptically to a new vial. An aliquot of 100 μl of each sample supernatant (containing the cell suspension) was mixed with an equal volume of 4 M sodium-azide solution to fix the cells. Fixed samples were kept at 4°C until flow cytometry counting (see below).

The rest of the supernatant cell suspension (~19 ml) was centrifuged in a swing-out rotor (Eppendorf A-4-62 Swing Bucket Rotor) at 3200 × *g* for 10 min to pellet cells. The liquid was discarded and cell pellets were frozen at −80°C until DNA isolation. Cell pellets were thawed and DNA was purified using a DNeasy PowerSoil kit (Qiagen) according to manufacturer’s protocol. The concentration of purified DNA was measured using a Qubit dsDNA BR Assay Kit (Invitrogen). DNA samples were stored at −20°C until library preparation (see below).

### Flow cytometry

Cell suspensions were filtered using a 40–μm nylon cell strainer (Falcon) and then fixed (see above). Fixed cell suspensions were serially diluted in sterile saline and stained with SYBR Green I for 15 min in the dark according to instructions of the supplier (Invitrogen). Stained cells suspensions were counted in 20 μl sample volume at medium flow rate (60 μl min^−1^) using an ACEA NovoCyte Green flow cytometer (OMNI Life Science Agilent). The SYBR Green I signal was measured in the FITC-channels of the instrument. Based on buffer controls, events with FSC-H-values above 50 and FITC-H above 350 were considered to potentially originate from microbial cells. Uninoculated microcosms, extracted and fixed in the same way, served to quantify cell-free (e.g., colloidal particles) background, which was subtracted from inoculated microcosm samples.

### Identification of soil isolates

Each soil isolate was identified based on the near-full length 16S rRNA gene, amplified by PCR with Phusion U Hot Start PCR MasterMix (Thermo-Fischer Scientific) in presence of 0.5 mM betaine (Sigma-Aldrich) using universal bacterial primers (27F 5’ AGAGTTTGATCCTGGCTCAG and 1492R 5’ GGTTACCTTGTTACGACTT, or 27F_deg 5’ AGRGTTYGATYMTGGCTCAG and 1391R_v18 5’ GACGGGCGGTGWGTRCA) (*63*). Amplified DNA was purified using Gel and PCR Clean-up kits (Macherey-Nagel) and single-end Sanger-sequenced with the corresponding forward primer at Eurofins Scientific. Sequences were compared to the SILVA database (version 132) using Blast (*64*) with default parameters for the genus level identification.

### Community 16S rRNA gene amplicon sequencing

Aliquots of 10 ng purified DNA per sample were used to amplify the V3-V4 region of the bacterial 16S rRNA gene, following the Illumina 16S Metagenomic Sequencing Library protocol (https://support.illumina.com/documents/documentation/chemistry_documentation/16s/16s-metagenomic-library-prep-guide-15044223b.pdf), indexed with a set A Nextera XT Index Kit (v2, Illumina), quantified and pooled in equal amounts for sequencing. The pooled SynCom amplicon libraries were spiked with 25% PhiX control DNA and paired-end sequenced on an Illumina MiniSeq instrument with the mid–output flow cell (Illumina). NatCom libraries and a sample of the SynCom starting inoculum were sequenced on a MiSeq platform with 300 cycles MiSeq v3 paired-end sequencing at the Lausanne Genomic Technologies Facility. Given their known reduced composition, for SynCom samples only the V4-end reads were used for analysis. Raw sequence reads were quality checked using FastQC 0.11.7 (*65*), cleaned and trimmed where necessary using Trimmomatic 0.36 (*66*). Primer sequences, ends with low quality and reads with poor quality score were removed. The quality was re-checked after trimming. A reference database of the inoculated SynCom members was created using the determined 16S rRNA gene sequences of each isolate (described above) and complemented by all unique sequence variants obtained from a MiSeq paired-end analysis of the SynCom inoculum. These reads were processed with QIIME 2 on a Unix platform (version qiime2-2018.8) (*67*), and grouped into taxonomic units at level 6 at 99% sequence identity by comparison to the SILVA database (version 132). Sequences were aligned using MUSCLE 3.8.1551 (*68*) and visualized using Jalview (*69*). Unique variable regions of 60 or 90 bp length were selected as identifier for each of the 21 SynCom strains. Strain abundances in the SynCom samples were then counted in the pools of quality-controlled sequence reads by searching for the unique selected sequence identifiers of each member in the reference database, using the bash command “grep”. The obtained counts were corrected for the number of 16S rRNA operons in the respective SynCom isolates genomes (to be described elsewhere). Relative abundances were then normalized to the total number of classified reads in each sample, which was further compared to differences in total cell count (as determined by flow cytometry) and the concentration of purified sample DNA.

### Microbe Atlas comparison

All sample sequences were compared to a global background of soil communities from the Microbe Atlas Project database (MAPdb, https://microbeatlas.org). The raw 16S reads from all samples were standardized and quality-filtered using a custom C++ program employed internally by MAPdb and then mapped using MAPseq 1.2.6 (*70*) (reference database: MAPref v2.2; all other parameters kept at default) to obtain 97%-level OTU count tables compatible with MAPdb. Samples from MAPdb with meta-data annotations “soil” (main environment) or “rhizosphere” (sub-environment) were used for downstream analysis (110,928 samples total). Earth Microbiome Project (*71*) samples were identified based on accessions from https://ebi-metagenomics.github.io/blog/2019/04/17/Earth-Microbiome-Project/ and corresponding soil pH values were extracted via the “sample_ph” field from accession-matched Sequence Read Archive (*72*) annotation files.

### Simulation model

To test the effects of stochastic variations in starting numbers of rapidly growing members within complex communities, we deployed a recently developed community model that simulates substrate-limited Monod growth of large numbers of bacterial taxa simultaneously (*73*). The model was seeded with 200,000 individual cells sampled with a weighted probability distribution from the measured relative abundances of 314 major taxa in a soil sample. Growth rates were attributed between 0.01 and 0.4 h^−1^ according to the log10 relative taxa abundance at start, except for five taxa with subsampled starting numbers between 0 and 10 (of 200,000 cells in total) that were given growth rates of 0.55, 0.25, 0.8, 0.6 and 0.35 h^−1^. Growth was allowed to proceed until all carbon was depleted, after which the final community was subsampled to 200,000 cells (to resemble a sequenced sample with 2×10^5^ reads). Relative and stacked taxa abundances were plotted within these subsampled data sets. Simulations were repeated five times independently.

### Statistical analyses

Data were analysed using R 3.6.1 (R Core Team, 2019) and the R packages *vegan* (*74*), *ggplot2* (*75*), *phyloseq* (*76*), *reshape* (*77*) and also using GraphPad Prism (version 9.0.0 for Mac OS X). The trends of microcosm total cell densities (as measured by flow cytometry) were compared using ANCOVA (*n* = 4-10 replicates per condition). Absolute abundances per SynCom community member were calculated from their relative (sequence) abundance times the measured total community size per replicate (from flow cytometry). The influence of culturing environment (e.g., soil, liquid) on community yield was compared using one-tailed t-tests. Differences in DNA yields were compared using a one-way ANOVA with post-hoc Tukey’s multiple comparisons test. The inter-replicate variability was expressed by the Bray-Curtis replicate distance from the community centroid. Effects of conditions were compared using one-way ANOVA with post-hoc Tukey’s multiple comparisons test. Alpha diversity was computed as community richness and Shannon indices.

Communities at different time points and treatments were compared by non-metric multidimensional scaling (NMDS) using Bray-Curtis distance values of normalized relative community member abundances. Multivariate dispersion of the data was examined using the *betadisper* function from *vegan*. Adonis (MANOVA with 999 permutations) was used to assess the differences between groups based on the output of *vegdist* (Bray-Curtis distances). The effect of toluene exposure on community cell densities was assessed using a Wilcoxon matched-pairs signed rank test. The effect of time and toluene exposure on community richness and Shannon values was assessed using two-way ANOVA. Clustered heatmap and UMAP (*78*) projections were generated from Bray-Curtis distance matrices using julia 1.6.0 (*79*) and the *Distances.jl* package ((*80*), version 0.10.3). UMAP projections were computed using the *UMAP.jl* package (https://github.com/dillondaudert/UMAP.jl, version 0.1.8; parameters: n_neighbors=500, min_dist=1.5, spread=15, epochs=2000). Scatter plots were produced using python 3.9.1 (*81*) and the *seaborn* package ((*82*), version 0.11.0).

### Database access

The NatCom and SynCom sequencing data are available from the Short Read Archives under BioProject number PRJNA767350.

## Acknowledgements

The authors acknowledge the support of the Lausanne Genomics Technology Facility for the amplicon sequencing, and Michael Rowley, Thibault Lambert and Thierry Adatte from the Faculty of Geosciences and Environment, University of Lausanne, for their help in soil analysis.

## Author contributions

## REFERENCES

1. N. Fierer, Embracing the unknown: disentangling the complexities of the soil microbiome. Nat Rev Microbiol 15, 579 (2017).10.1038/nrmicro.2017.87

2. M. Bahram, F. Hildebrand, S. K. Forslund, J. L. Anderson, N. A. Soudzilovskaia, P. M. Bodegom, J. Bengtsson-Palme, S. Anslan, L. P. Coelho, H. Harend, J. Huerta-Cepas, M. H. Medema, M. R. Maltz, S. Mundra, P. A. Olsson, M. Pent, S. Põlme, S. Sunagawa, M. Ryberg, L. Tedersoo, P. Bork, Structure and function of the global topsoil microbiome. Nature 560, 233 (2018). 10.1038/s41586-018-0386-6

3. M. A. Moran, The global ocean microbiome. Science 350, aac8455 (2015).10.1126/science.aac8455

4. D. B. Müller, C. Vogel, Y. Bai, J. A. Vorholt, The plant microbiota: Systems-level insights and perspectives. Annu Rev Genet 50, 211 (2016).10.1146/annurev-genet-120215-034952

5. D. Berry, A. Loy, Stable-isotope probing of human and animal microbiome function. Trends Microbiol 26, 999 (2018).https://doi.org/10.1016/j.tim.2018.06.004

6. P. Engel, W. K. Kwong, Q. McFrederick, K. E. Anderson, S. M. Barribeau, J. A. Chandler, R. S. Cornman, J. Dainat, J. R. d. Miranda, V. Doublet, O. Emery, J. D. Evans, L. Farinelli, M. L. Flenniken, F. Granberg, J. A. Grasis, L. Gauthier, J. Hayer, H. Koch, S. Kocher, V. G. Martinson, N. Moran, M. Munoz-Torres, I. Newton, R. J. Paxton, E. Powell, B. M. Sadd, P. Schmid-Hempel, R. Schmid-Hempel, S. J. Song, R. S. Schwarz, D. vanEngelsdorp, B. Dainat, G. B. Hurst, R. J. Collier, The bee microbiome: Impact on bee health and model for evolution and ecology of host-microbe interactions. mBio 7, e02164 (2016).doi:10.1128/mBio.02164-15

7. K. Hosoda, S. Suzuki, Y. Yamauchi, Y. Shiroguchi, A. Kashiwagi, N. Ono, K. Mori, T. Yomo, Cooperative adaptation to establishment of a synthetic bacterial mutualism. PLoS One 6, e17105 (2011).10.1371/journal.pone.0017105

8. Y. Tanouchi, R. P. Smith, L. You, Engineering microbial systems to explore ecological and evolutionary dynamics. Curr Opin Biotechnol 23, 791 (2012).10.1016/j.copbio.2012.01.006

9. H. Celiker, J. Gore, Clustering in community structure across replicate ecosystems following a long-term bacterial evolution experiment. Nat Commun 5, 4643 (2014).10.1038/ncomms5643

10. J. Friedman, L. M. Higgins, J. Gore, Community structure follows simple assembly rules in microbial microcosms. Nat Ecol Evol 1, 109 (2017).10.1038/s41559-017-0109

11. S. G. Hays, W. G. Patrick, M. Ziesack, N. Oxman, P. A. Silver, Better together: engineering and application of microbial symbioses. Curr Opin Biotechnol 36, 40 (2015).10.1016/j.copbio.2015.08.008

12. B. E. L. Morris, R. Henneberger, H. Huber, C. Moissl-Eichinger, Microbial syntrophy: interaction for the common good. FEMS Microbiol Rev 37, 384 (2013).10.1111/1574-6976.12019

13. B. Borer, D. Ciccarese, D. Johnson, D. Or, Spatial organization in microbial range expansion emerges from trophic dependencies and successful lineages. Comm Biol 3, 685 (2020).10.1038/s42003-020-01409-y

14. E. E. Lilja, D. R. Johnson, Segregating metabolic processes into different microbial cells accelerates the consumption of inhibitory substrates. ISME J 10, 1568 (2016).10.1038/ismej.2015.243

15. E. H. Wintermute, P. A. Silver, Dynamics in the mixed microbial concourse. Genes Dev 24, 2603 (2010).10.1101/gad.1985210

16. T. A. Hoek, K. Axelrod, T. Biancalani, E. A. Yurtsev, J. Liu, J. Gore, Resource availability modulates the cooperative and competitive nature of a microbial cross-feeding mutualism. PLoS Biol 14, e1002540 (2016).10.1371/journal.pbio.1002540

17. M. T. Mee, J. J. Collins, G. M. Church, H. H. Wang, Syntrophic exchange in synthetic microbial communities. Proc Natl Acad Sci U S A 111, E2149 (2014).10.1073/pnas.1405641111

18. F. Goldschmidt, R. R. Regoes, D. R. Johnson, Successive range expansion promotes diversity and accelerates evolution in spatially structured microbial populations. ISME J 11, 2112 (2017).10.1038/ismej.2017.76

19. R. H. Hsu, R. L. Clark, J. W. Tan, J. C. Ahn, S. Gupta, P. A. Romero, O. S. Venturelli, Microbial interaction network inference in microfluidic droplets. Cell Syst, (2019).10.1016/j.cels.2019.06.008

20. M. Manhart, E. I. Shakhnovich, Growth tradeoffs produce complex microbial communities on a single limiting resource. Nat Commun 9, 3214 (2018).10.1038/s41467-018-05703-6

21. L. S. Bittleston, M. Gralka, G. E. Leventhal, I. Mizrahi, O. X. Cordero, Context-dependent dynamics lead to the assembly of functionally distinct microbial communities. Nat Commun 11, 1440 (2020).10.1038/s41467-020-15169-0

22. E. S. Wright, R. Gupta, K. H. Vetsigian, Multi-stable bacterial communities exhibit extreme sensitivity to initial conditions. FEMS Microbiol Ecol 97, (2021).10.1093/femsec/fiab073

23. A. R. Pacheco, M. L. Osborne, D. Segrè, Non-additive microbial community responses to environmental complexity. Nat Commun 12, 2365 (2021).10.1038/s41467-021-22426-3

24. A. Dal Co, S. van Vliet, D. J. Kiviet, S. Schlegel, M. Ackermann, Short-range interactions govern the dynamics and functions of microbial communities. Nat Ecol Evol 4, 366 (2020).10.1038/s41559-019-1080-2

25. M. Dubey, N. Hadadi, S. Pelet, N. Carraro, D. R. Johnson, J. R. van der Meer, Environmental connectivity controls diversity in soil microbial communities. Comm Biol 4, 492 (2021).10.1038/s42003-021-02023-2

26. B. Kerr, M. A. Riley, M. W. Feldman, B. J. M. Bohannan, Local dispersal promotes biodiversity in a real-life game of rock–paper–scissors. Nature 418, 171 (2002).10.1038/nature00823

27. E. D. Kelsic, J. Zhao, K. Vetsigian, R. Kishony, Counteraction of antibiotic production and degradation stabilizes microbial communities. Nature 521, 516 (2015).10.1038/nature14485

28. H. J. Kim, J. Q. Boedicker, J. W. Choi, R. F. Ismagilov, Defined spatial structure stabilizes a synthetic multispecies bacterial community. Proc Natl Acad Sci U S A 105, 18188 (2008).10.1073/pnas.0807935105

29. L. F. W. Roesch, R. R. Fulthorpe, A. Riva, G. Casella, A. K. M. Hadwin, A. D. Kent, S. H. Daroub, F. A. O. Camargo, W. G. Farmerie, E. W. Triplett, Pyrosequencing enumerates and contrasts soil microbial diversity. ISME J 1, 283 (2007).10.1038/ismej.2007.53

30. X. Raynaud, N. Nunan, Spatial ecology of bacteria at the microscale in soil. PLoS One 9, e87217 (2014).10.1371/journal.pone.0087217

31. J. K. Jansson, K. S. Hofmockel, The soil microbiome—from metagenomics to metaphenomics. Curr Opin Microbiol 43, 162 (2018).https://doi.org/10.1016/j.mib.2018.01.013

32. M. G. A. van der Heijden, R. D. Bardgett, N. M. van Straalen, The unseen majority: soil microbes as drivers of plant diversity and productivity in terrestrial ecosystems. Ecol Lett 11, 296 (2008).10.1111/j.1461-0248.2007.01139.x

33. R. Amundson, A. A. Berhe, J. W. Hopmans, C. Olson, A. E. Sztein, D. L. Sparks, Soil and human security in the 21st century. Science 348, 1261071 (2015).10.1126/science.1261071

34. J. Kaiser, Wounding Earth’s fragile skin. Science 304, 1616 (2004).10.1126/science.304.5677.1616

35. G. K. Mishra, Microbes in heavy metal remediation: A review on current trends and patents. Recent Pat Biotechnol 11, 188 (2017).10.2174/1872208311666170120121025

36. T. M. Vogel, Bioaugmentation as a soil bioremediation approach. Curr Opin Biotechnol 7, 311 (1996).https://doi.org/10.1016/S0958-1669(96)80036-X

37. A. Mrozik, Z. Piotrowska-Seget, Bioaugmentation as a strategy for cleaning up of soils contaminated with aromatic compounds. Microbiol Res 165, 363 (2010).https://doi.org/10.1016/j.micres.2009.08.001

38. P. H. Pritchard, Use of inoculation in bioremediation. Curr Opin Biotechnol 3, 232 (1992).https://doi.org/10.1016/0958-1669(92)90098-4

39. J. B. Fellman, E. Hood, R. G. M. Spencer, Fluorescence spectroscopy opens new windows into dissolved organic matter dynamics in freshwater ecosystems: A review. Limnol Oceanogr 55, 2452 (2010).10.4319/lo.2010.55.6.2452

40. B. Gworek, A. H. Baczewska-Dabrowska, R. Kalinowski, E. B. Gorska, H. Rekosz-Burlaga, D. Gozdowski, I. Olejniczak, M. Graniewska, W. Dmuchowski, Ecological risk assessment for land contaminated by petrochemical industry. PLoS One 13, e0204852 (2018).10.1371/journal.pone.0204852

41. J. A. Vorholt, C. Vogel, C. I. Carlström, D. B. Müller, Establishing causality: Opportunities of synthetic communities for plant microbiome research. Cell Host & Microbe 22, 142 (2017).https://doi.org/10.1016/j.chom.2017.07.004

42. A. Shade, H. Peter, S. Allison, D. Baho, M. Berga, H. Buergmann, D. Huber, S. Langenheder, J. Lennon, J. Martiny, K. Matulich, T. Schmidt, J. Handelsman, Fundamentals of microbial community resistance and resilience. Front Microbiol 3, (2012).10.3389/fmicb.2012.00417

43. C. S. Holling, Resilience and stability of ecological systems. Annu Rev Ecol System 4, 1 (1973).10.1146/annurev.es.04.110173.000245

44. B. Niu, J. N. Paulson, X. Zheng, R. Kolter, Simplified and representative bacterial community of maize roots. Proc Natl Acad Sci U S A 114, E2450 (2017).10.1073/pnas.1616148114

45. D. R. Nemergut, S. K. Schmidt, T. Fukami, S. P. O’Neill, T. M. Bilinski, L. F. Stanish, J. E. Knelman, J. L. Darcy, R. C. Lynch, P. Wickey, S. Ferrenberg, Patterns and processes of microbial community assembly. Microbiol Mol Biol Rev 77, 342 (2013).doi:10.1128/MMBR.00051-12

46. S. Widder, R. J. Allen, T. Pfeiffer, T. P. Curtis, C. Wiuf, W. T. Sloan, O. X. Cordero, S. P. Brown, B. Momeni, W. Shou, H. Kettle, H. J. Flint, A. F. Haas, B. Laroche, J.-U. Kreft, P. B. Rainey, S. Freilich, S. Schuster, K. Milferstedt, J. R. van der Meer, T. Groβkopf, J. Huisman, A. Free, C. Picioreanu, C. Quince, I. Klapper, S. Labarthe, B. F. Smets, H. Wang, O. S. Soyer, F. Isaac Newton Institute, Challenges in microbial ecology: building predictive understanding of community function and dynamics. ISME J 10, 2557 (2016).10.1038/ismej.2016.45

47. D. Naylor, S. Fansler, C. Brislawn, W. C. Nelson, K. S. Hofmockel, J. K. Jansson, R. McClure, Deconstructing the soil microbiome into reduced-complexity functional modules. mBio 11, e01349 (2020).10.1128/mBio.01349-20

48. E. M. Bach, R. J. Williams, S. K. Hargreaves, F. Yang, K. S. Hofmockel, Greatest soil microbial diversity found in micro-habitats. Soil Biol Biochem 118, 217 (2018).https://doi.org/10.1016/j.soilbio.2017.12.018

49. J. E. Goldford, N. Lu, D. Bajić, S. Estrela, M. Tikhonov, A. Sanchez-Gorostiaga, D. Segrè, P. Mehta, A. Sanchez, Emergent simplicity in microbial community assembly. Science (New York, N.Y.) 361, 469 (2018).10.1126/science.aat1168

50. R. L. Wilpiszeski, J. A. Aufrecht, S. T. Retterer, M. B. Sullivan, D. E. Graham, E. M. Pierce, O. D. Zablocki, A. V. Palumbo, D. A. Elias, Soil aggregate microbial communities: Towards understanding microbiome interactions at biologically relevant scales. Applied and environmental microbiology 85, e00324 (2019).10.1128/AEM.00324-19

51. O. X. Cordero, M. S. Datta, Microbial interactions and community assembly at microscales. Curr Opin Microbiol 31, 227 (2016).https://doi.org/10.1016/j.mib.2016.03.015

52. G. W. Nicol, D. Tscherko, T. M. Embley, J. I. Prosser, Primary succession of soil Crenarchaeota across a receding glacier foreland. Environ Microbiol 7, 337 (2005).10.1111/j.1462-2920.2005.00698.x

53. D. Górniak, H. Marszałek, M. Kwaśniak-Kominek, G. Rzepa, M. Manecki, Soil formation and initial microbiological activity on a foreland of an Arctic glacier (SW Svalbard). Appl Soil Ecol 114, 34 (2017).https://doi.org/10.1016/j.apsoil.2017.02.017

54. M. Kim, D. Or, Hydration status and diurnal trophic interactions shape microbial community function in desert biocrusts. Biogeosciences 14, 5403 (2017).10.5194/bg-14-5403-2017

55. S. Schmidt, E. Costello, D. Nemergut, C. C. Cleveland, S. Reed, M. Weintraub, A. Meyer, A. Martin, Biogeochemical consequences of rapid microbial turnover and seasonal succession in soil. Ecology 88, 1379 (2007)

56. S. C. Neubauer, K. Givler, S. Valentine, J. P. Megonigal, Seasonal patterns and plant-mediated controls of subsurface wetland biogeochemistry. Ecology 86, 3334 (2005)

57. A. E. Magurran, P. A. Henderson, Explaining the excess of rare species in natural species abundance distributions. Nature 422, 714 (2003).10.1038/nature01547

58. M. L. Sogin, H. G. Morrison, J. A. Huber, D. Mark Welch, S. M. Huse, P. R. Neal, J. M. Arrieta, G. J. Herndl, Microbial diversity in the deep sea and the underexplored “rare biosphere”. Proc Natl Acad Sci U S A 103, 12115 (2006).10.1073/pnas.0605127103

59. V. Kurm, W. H. van der Putten, W. de Boer, S. Naus-Wiezer, W. H. Hol, Low abundant soil bacteria can be metabolically versatile and fast growing. Ecology 98, 555 (2017).10.1002/ecy.1670

60. A. Shade, S. E. Jones, J. G. Caporaso, J. Handelsman, R. Knight, N. Fierer, J. A. Gilbert, N. Dubilier, Conditionally rare taxa disproportionately contribute to temporal changes in microbial diversity. mBio 5, e01371 (2014).doi:10.1128/mBio.01371-14

61. J. Liu, J. Li, L. Feng, H. Cao, Z. Cui, An improved method for extracting bacteria from soil for high molecular weight DNA recovery and BAC library construction. J Microbiol 48, 728 (2010).10.1007/s12275-010-0139-1

62. D. L. Balkwill, W. C. Ghiorse, Characterization of subsurface bacteria associated with two shallow aquifers in Oklahoma. Applied and environmental microbiology 50, 580 (1985)

63. D. J. Lane, B. Pace, G. J. Olsen, D. A. Stahl, M. L. Sogin, N. R. Pace, Rapid determination of 16S ribosomal RNA sequences for phylogenetic analyses. Proc Natl Acad Sci U S A 82, 6955 (1985).10.1073/pnas.82.20.6955

64. E. Pruesse, J. Peplies, F. O. Glockner, SINA: accurate high-throughput multiple sequence alignment of ribosomal RNA genes. Bioinformatics 28, 1823 (2012).10.1093/bioinformatics/bts252

65. S. Andrews, FastQC: A quality control tool for high throughput sequence data.

66. A. M. Bolger, M. Lohse, B. Usadel, Trimmomatic: a flexible trimmer for Illumina sequence data. Bioinformatics 30, 2114 (2014).10.1093/bioinformatics/btu170

67. E. Bolyen, J. R. Rideout, M. R. Dillon, N. A. Bokulich, C. C. Abnet, G. A. Al-Ghalith, H. Alexander, E. J. Alm, M. Arumugam, F. Asnicar, Y. Bai, J. E. Bisanz, K. Bittinger, A. Brejnrod, C. J. Brislawn, C. T. Brown, B. J. Callahan, A. M. Caraballo-Rodriguez, J. Chase, E. K. Cope, R. Da Silva, C. Diener, P. C. Dorrestein, G. M. Douglas, D. M. Durall, C. Duvallet, C. F. Edwardson, M. Ernst, M. Estaki, J. Fouquier, J. M. Gauglitz, S. M. Gibbons, D. L. Gibson, A. Gonzalez, K. Gorlick, J. Guo, B. Hillmann, S. Holmes, H. Holste, C. Huttenhower, G. A. Huttley, S. Janssen, A. K. Jarmusch, L. Jiang, B. D. Kaehler, K. B. Kang, C. R. Keefe, P. Keim, S. T. Kelley, D. Knights, I. Koester, T. Kosciolek, J. Kreps, M. G. I. Langille, J. Lee, R. Ley, Y. X. Liu, E. Loftfield, C. Lozupone, M. Maher, C. Marotz, B. D. Martin, D. McDonald, L. J. McIver, A. V. Melnik, J. L. Metcalf, S. C. Morgan, J. T. Morton, A. T. Naimey, J. A. Navas-Molina, L. F. Nothias, S. B. Orchanian, T. Pearson, S. L. Peoples, D. Petras, M. L. Preuss, E. Pruesse, L. B. Rasmussen, A. Rivers, M. S. Robeson, 2nd, P. Rosenthal, N. Segata, M. Shaffer, A. Shiffer, R. Sinha, S. J. Song, J. R. Spear, A. D. Swafford, L. R. Thompson, P. J. Torres, P. Trinh, A. Tripathi, P. J. Turnbaugh, S. Ul-Hasan, J. J. J. van der Hooft, F. Vargas, Y. Vazquez-Baeza, E. Vogtmann, M. von Hippel, W. Walters, Y. Wan, M. Wang, J. Warren, K. C. Weber, C. H. D. Williamson, A. D. Willis, Z. Z. Xu, J. R. Zaneveld, Y. Zhang, Q. Zhu, R. Knight, J. G. Caporaso, Reproducible, interactive, scalable and extensible microbiome data science using QIIME 2. Nat Biotechnol 37, 852 (2019).10.1038/s41587-019-0209-9

68. R. C. Edgar, MUSCLE: multiple sequence alignment with high accuracy and high throughput. Nucleic Acids Res 32, 1792 (2004).10.1093/nar/gkh340

69. A. M. Waterhouse, J. B. Procter, D. M. Martin, M. Clamp, G. J. Barton, Jalview Version 2--a multiple sequence alignment editor and analysis workbench. Bioinformatics 25, 1189 (2009).10.1093/bioinformatics/btp033

70. J. F. Matias Rodrigues, T. S. B. Schmidt, J. Tackmann, C. von Mering, MAPseq: highly efficient k-mer search with confidence estimates, for rRNA sequence analysis. Bioinformatics 33, 3808 (2017).10.1093/bioinformatics/btx517

71. L. R. Thompson, J. G. Sanders, D. McDonald, A. Amir, J. Ladau, K. J. Locey, R. J. Prill, A. Tripathi, S. M. Gibbons, G. Ackermann, J. A. Navas-Molina, S. Janssen, E. Kopylova, Y. Vazquez-Baeza, A. Gonzalez, J. T. Morton, S. Mirarab, Z. Zech Xu, L. Jiang, M. F. Haroon, J. Kanbar, Q. Zhu, S. Jin Song, T. Kosciolek, N. A. Bokulich, J. Lefler, C. J. Brislawn, G. Humphrey, S. M. Owens, J. Hampton-Marcell, D. Berg-Lyons, V. McKenzie, N. Fierer, J. A. Fuhrman, A. Clauset, R. L. Stevens, A. Shade, K. S. Pollard, K. D. Goodwin, J. K. Jansson, J. A. Gilbert, R. Knight, C. Earth Microbiome Project, A communal catalogue reveals Earth’s multiscale microbial diversity. Nature 551, 457 (2017).10.1038/nature24621

72. R. Leinonen, H. Sugawara, M. Shumway, I. N. S. D. Collaboration., The sequence read archive. Nucleic Acids Res 39 D19 (2010)

73. N. Hadadi, J. R. van der Meer. (Zenodo, 2021). http://doi.org/10.5281/zenodo.4568347

74. J. Oksanen, R. Kindt, P. Legendre, B. O’Hara, M. H. H. Stevens, M. J. Oksanen, M. Suggests, The vegan package. Community ecology package 10, 719 (2007)

75. H. Wickham, ggplot2. Wiley Interdisciplinary Reviews: Computational Statistics 3, 180 (2011)

76. P. J. McMurdie, S. Holmes, phyloseq: an R package for reproducible interactive analysis and graphics of microbiome census data. PLoS one 8, e61217 (2013)

77. H. Wickham, Reshaping data with the reshape package. J Stat Software 21, 1 (2007)

78. E. Becht, L. McInnes, J. Healy, C. A. Dutertre, I. W. H. Kwok, L. G. Ng, F. Ginhoux, E. W. Newell, Dimensionality reduction for visualizing single-cell data using UMAP. Nat Biotechnol 37, 38 (2019).10.1038/nbt.4314

79. J. Bezanson, A. Edelman, S. Karpinski, V. B. Shah, Julia: A fresh approach to numerical computing. SIAM Rev. 59, 65 (2017).doi:10.1137/141000671.

80. JuliaStats, Distances.jl, a Julia package for evaluating distances (metrics) between vectors.

81. G. Van Rossum, F. L. Drake, (CreateSpace., Scotts Valley, CA, 2009).

82. M. L. Waskom, Seaborn: statistical data visualization. J Open Source Software 6, 3021 (2021)

