## Supplementary Methods Tables S1-S4 Figures S1-S6 for "Reproducible Propagation of Species-Rich Soil Microbiomes Suggests Robust Underlying Deterministic Principles of Community Formation"

**To**

**Supplementary Methods**
**Tables S1-S4**
**Figures S1-S6**
**Supplementary References**

**Supplementary Methods**

**Analysis of soil parameters.** The gravimetric water content in twice autoclaved soil was determined from weight loss of soil samples before and after drying at 70°C for 10 days. Soil-pH was measured in mixed solution with distilled water, stirred for one hour at 120 rpm, using an Orion Star A111 Benchtop pH Meter (Thermo Fisher Scientific).

Organic material was characterized by UV/Vis and fluorescence spectrometry. Soil-water (5 g) or SE-water (5 ml) extracts were prepared by mixing sample in 12 replicates with 20 ml MilliQ water at 80 rpm for 1 h (Supplementary table S1). Mixtures were subsequently centrifuged for 15 min at 4600 × g and the supernatant was filtered using a 0.2–µm Stericup Quick Release System PES filter (Merck). Filtered samples were stored in glass amber vials at 4˚C in the dark prior to analysis.

Filtered samples were serially diluted in MilliQ water, transferred to 1–cm quartz cuvettes and measured in a UV/Vis spectrophotometer (Perkin Elmer 650S) or a Fluorolog-3 spectrofluorometer (Horiba). Data were collected in the three-dimensional form of excitation-emission matrices (EEMs) for a parallel factor analysis (PARAFAC) model, against MilliQ water. Excitation wavelengths ranged from 270 to 500 nm and emissions were measured in the range from 300 to 600 nm. Data were processed using the PARAFAC algorithms (*1*) in MATLAB (vs.2016a, MathWorks). Detected spectra correspond to six different organic matter types as described by Fellman et al., 2010 (*2*) (Supplementary table S2). $NH_4$-N, $NO_3$-N and total-N in the final soil+SE was determined by Sol-Conseil (Gland, Switzerland).

**RockEval methodology.** RockEval analysis was used to assess the carbon content composition of natural soil, autoclaved soil and soil mixed with SE, as suggested (*3*). Upon mixing and drying to remove the remaining water content, the samples were grounded using a Planetary Micro Mili Pulverisette 7 (Fritsch). The samples (including the IFP160000 standard) were processed using a RockEval 6 Pyrolyser (Vinci Technologies) at the Faculty of Geosciences and Environment, University of Lausanne. In short, samples were pyrolysed and combusted, leading to the release of hydrocarbons ($S_1$ peak), kerogen

($S_2$), and $CO_2$ ($S_3$), and remainder residual carbon (RC), which were measured by flame ionization and thermal conductivity detectors (*3*). The obtained values of $S_1$, $S_2$ and $S_3$ were used to calculate the total organic carbon (TOC), pyrolyzable and mineral carbon fractions, and the hydrogen and oxygen indices (Supplementary table S3).

The working principle of the apparatus consists of pyrolysis followed by the combustion of the sample, which first leads to the release of hydrocarbons ($S_1$ peak), kerogen ($S_2$), and $CO_2$ ($S_3$). Residual carbon (RC) is also quantified. $T_{max}$ is the temperature needed for maximal generation of hydrocarbons and is indicative of thermal maturity. FID (flame ionization detector) and TCD (thermal conductivity detector) are detectors used in measurements. RockEval provides insight into: TOC (total organic carbon), PC (pyrolyzable carbon), MINC (mineral carbon), HI (hydrogen index) and OI (oxygen index) derived from the values of $S_1$, $S_2$ and $S_3$ and each other.

HI represents the ratio of hydrogen to organic carbon and is indicative of the origin of the organic material. OI shows the amount of oxygen relative to TOC. These indices are calculated as follows:

$HI=S_2/TOCx100$;

$OI=S_3/TOCx100$

TOC of the SE-solution was determined by Scitec Research SA (Lausanne, Switzerland).

**Table S1. UV-Vis fluorescence measurement values expressed as total fluorescence, Napierian absorbance and absorbance ratio 250:365.**

| | **Autoclaved soil** | **Soil** | **Soil extract** | **Autoclaved soil & extract** |
|---|---|---|---|---|
| **$F_{tot}$** | 5.08±0.24 | 3.14±0.04 | 127.26±5.71 | 11.61±0.11 |
| **Napierian absorbance** | 22.69±0.47 | 12.9±0.24 | 1413.7±90.96 | 62.32±1.42 |
| **A250:A365** | 4.47±0.08 | 4.55±0.09 | 5.28±0.17 | 4.79±0.1 |

**Table S2. Classification of PARAFAC components.**

| **Comp[a].** | **Max1 Ex (nm)** | **Max Em (nm)** | **Description** |
|---|---|---|---|
| **C1** | <270 (445) | 538 | UVA Humic-like (high molecular weight, aromatic, fluorescence resembles fulvic acid, widespread) |
| **C2** | 310 | 422 | UVA Humic-like (low molecular weight, common in marine environments associated with biological activity but can be found in wastewater, wetland, and agricultural environments) |
| **C3** | <270 (340) | 472 | UVC Humic-like (high molecular weight, aromatic, fl acid, widespread) |
| **C4** | 270 | 310 | Tyrosine-like (amino acids, free or bound in proteins, fluorescence resembles free tyrosine, may indicate more degraded peptide material) |
| **C5** | <270 | 440 | UVA Humic-like (high molecular weight and aromatic humic, widespread, but highest in wetlands and forested environments) |
| **C6** | 280 | 346 | Tryptophane-like (amino acids, free or bound in proteins, fluorescence resembles free tryptophan) |

a) classification based on Ref. (*2*)

**Table S3. Analysis of soil organic carbon and nitrogen**

| Sample | PC [%] | RC [%] | TOC [%] | MIN C [%] | HI [mg HC/g TOC] | OI [mg $CO_2$/g TOC] | $NO_3$-N [mg/kg] | $NH_4$-N [mg/kg] | Total-N [%] |
|---|---|---|---|---|---|---|---|---|---|
| Soil | 0.04 | 0.10 | 0.13 | 2.75 | 151 | 563 | nd | nd | nd |
| Autoclaved soil | 0.04 | 0.11 | 0.15 | 3.62 | 161 | 426 | nd | nd | nd |
| Autoclaved soil & SE | 0.03 | 0.12 | 0.15 | 3.06 | 116 | 379 | <0.01 | 2.31 | 0.03 |

**Table S4. List of all strains isolated from forest soil**

| No. | slv_last taxa | phyla;class |
|---|---|---|
| 1 | Actinobacteria | Proteobacteria;Gammaproteobacteria |
| 2 | Agromyces | Actinobacteria;Actinobacteria |
| 3 | Agromyces | Actinobacteria;Actinobacteria |
| 4 | Agromyces | Actinobacteria;Actinobacteria |
| 5 | Agromyces | Actinobacteria;Actinobacteria |
| 6 | Agromyces | Actinobacteria;Actinobacteria |
| 7 | Agromyces | Actinobacteria;Actinobacteria |
| 8 | Agromyces | Actinobacteria;Actinobacteria |
| 9 | Agromyces | Actinobacteria;Actinobacteria |
| 10 | Allorhizobium-Neorhizobium-Pararhizobium-Rhizobium | Proteobacteria;Alphaproteobacteria |
| 11 | Allorhizobium-Neorhizobium-Pararhizobium-Rhizobium | Proteobacteria;Alphaproteobacteria |
| 12 | Allorhizobium-Neorhizobium-Pararhizobium-Rhizobium; | Proteobacteria;Alphaproteobacteria |
| 13 | Altererythrobacter | Proteobacteria;Alphaproteobacteria |
| 14 | Aminobacter | Proteobacteria;Alphaproteobacteria |
| 15 | Angustibacter | Actinobacteria;Actinobacteria |
| 16 | Angustibacter | Actinobacteria;Actinobacteria |
| 17 | Angustibacter | Actinobacteria;Actinobacteria |
| 18 | Angustibacter | Actinobacteria;Actinobacteria |
| 19 | Angustibacter | Actinobacteria;Actinobacteria |
| 20 | Bosea | Proteobacteria;Alphaproteobacteria |
| 21 | Bosea | Proteobacteria;Alphaproteobacteria |
| 22 | Bosea | Proteobacteria;Alphaproteobacteria |
| 23 | Bradyrhizobium | Proteobacteria;Alphaproteobacteria |
| 24 | Bradyrhizobium | Proteobacteria;Alphaproteobacteria |
| 25 | Bradyrhizobium | Proteobacteria;Alphaproteobacteria |
| 26 | Bradyrhizobium | Proteobacteria;Alphaproteobacteria |
| 27 | Bradyrhizobium | Proteobacteria;Alphaproteobacteria |
| 28 | Bradyrhizobium | Proteobacteria;Alphaproteobacteria |
| 29 | Bradyrhizobium | Proteobacteria;Alphaproteobacteria |
| 30 | Bradyrhizobium | Proteobacteria;Alphaproteobacteria |
| 31 | Bradyrhizobium | Proteobacteria;Alphaproteobacteria |
| 32 | Bradyrhizobium | Proteobacteria;Alphaproteobacteria |
| 33 | Bradyrhizobium | Proteobacteria;Alphaproteobacteria |
| 34 | Bradyrhizobium | Proteobacteria;Alphaproteobacteria |
| 35 | Bradyrhizobium | Proteobacteria;Alphaproteobacteria |
| 36 | Bradyrhizobium | Proteobacteria;Alphaproteobacteria |
| 37 | Burkholderia-Caballeronia-Paraburkholderia | Proteobacteria;Gammaproteobacteria |
| 38 | Burkholderia-Caballeronia-Paraburkholderia | Proteobacteria;Betaproteobacteria |
| 39 | Burkholderia-Caballeronia-Paraburkholderia | Proteobacteria;Betaproteobacteria |
| 40 | Burkholderia-Caballeronia-Paraburkholderia | Proteobacteria;Betaproteobacteria |
| 41 | Burkholderia-Caballeronia-Paraburkholderia | Proteobacteria;Betaproteobacteria |
| 42 | Burkholderia-Caballeronia-Paraburkholderia | Proteobacteria;Betaproteobacteria |
| 43 | Burkholderia-Caballeronia-Paraburkholderia | Proteobacteria;Betaproteobacteria |

| | | |
|---|---|---|
| 44 | Burkholderia-Caballeronia-Paraburkholderia | Proteobacteria;Betaproteobacteria |
| 45 | Burkholderia-Caballeronia-Paraburkholderia | Proteobacteria;Betaproteobacteria |
| 46 | Burkholderia-Caballeronia-Paraburkholderia | Proteobacteria;Betaproteobacteria |
| 47 | Burkholderia-Caballeronia-Paraburkholderia | Proteobacteria;Betaproteobacteria |
| 48 | Burkholderia-Caballeronia-Paraburkholderia | Proteobacteria;Betaproteobacteria |
| 49 | Burkholderia-Caballeronia-Paraburkholderia | Proteobacteria;Betaproteobacteria |
| 50 | Burkholderia-Caballeronia-Paraburkholderia | Proteobacteria;Betaproteobacteria |
| 51 | Burkholderiaceae | Proteobacteria;Gammaproteobacteria |
| 52 | Burkholderiaceae | Proteobacteria;Gammaproteobacteria |
| 53 | Caulobacter | Proteobacteria;Alphaproteobacteria |
| 54 | Caulobacter | Proteobacteria;Alphaproteobacteria |
| 55 | Caulobacter | Proteobacteria;Alphaproteobacteria |
| 56 | Caulobacter | Proteobacteria;Alphaproteobacteria |
| 57 | Caulobacter | Proteobacteria;Alphaproteobacteria |
| 58 | Cellulomonas | Actinobacteria;Actinobacteria |
| 59 | Cellulomonas | Actinobacteria;Actinobacteria |
| 60 | Cellulomonas | Actinobacteria;Actinobacteria |
| 61 | Cellulomonas | Actinobacteria;Actinobacteria |
| 62 | Cellulomonas | Actinobacteria;Actinobacteria |
| 63 | Chitinophaga | Bacteroidetes;Bacteroidia |
| 64 | Chitinophaga | Bacteroidetes;Bacteroidia |
| 65 | Cohnella | Firmicutes;Bacilli |
| 66 | Curtobacterium | Actinobacteria;Actinobacteria |
| 67 | Curtobacterium | Actinobacteria;Actinobacteria |
| 68 | Devosia | Proteobacteria;Alphaproteobacteria |
| 69 | Dyella | Proteobacteria;Gammaproteobacteria |
| 70 | Dyella | Proteobacteria;Gammaproteobacteria |
| 71 | Ensifer | Proteobacteria;Alphaproteobacteria |
| 72 | Ensifer | Proteobacteria;Alphaproteobacteria |
| 73 | Enterobacteriaceae | Proteobacteria;Gammaproteobacteria |
| 74 | Enterobacteriaceae | Proteobacteria;Gammaproteobacteria |
| 75 | Enterobacteriaceae | Proteobacteria;Gammaproteobacteria |
| 76 | Enterobacteriaceae | Proteobacteria;Gammaproteobacteria |
| 77 | Enterobacteriaceae | Proteobacteria;Gammaproteobacteria |
| 78 | Enterobacteriaceae | Proteobacteria;Gammaproteobacteria |
| 79 | Flavobacterium | Bacteroidetes;Bacteroidia |
| 80 | Flavobacterium | Bacteroidetes;Bacteroidia |
| 81 | Flavobacterium | Bacteroidetes;Bacteroidia |
| 82 | Frondihabitans | Actinobacteria;Actinobacteria |
| 83 | Frondihabitans | Actinobacteria;Actinobacteria |
| 84 | Kluyvera | Proteobacteria;Gammaproteobacteria |
| 85 | Labrys | Proteobacteria;Alphaproteobacteria |
| 86 | Labrys | Proteobacteria;Alphaproteobacteria |
| 87 | Labrys | Proteobacteria;Alphaproteobacteria |
| 88 | Leifsonia | Actinobacteria;Actinobacteria |
| 89 | Leifsonia | Actinobacteria;Actinobacteria |
| 90 | Luteibacter | Proteobacteria;Gammaproteobacteria |
| 91 | Luteibacter | Proteobacteria;Gammaproteobacteria |
| 92 | Lysobacter | Proteobacteria;Gammaproteobacteria |

| | | |
|---|---|---|
| 93 | Mesorhizobium | Proteobacteria;Alphaproteobacteria |
| 94 | Mesorhizobium | Proteobacteria;Alphaproteobacteria |
| 95 | Mesorhizobium | Proteobacteria;Alphaproteobacteria |
| 96 | Mesorhizobium | Proteobacteria;Alphaproteobacteria |
| 97 | Mesorhizobium | Proteobacteria;Alphaproteobacteria |
| 98 | Mesorhizobium | Proteobacteria;Alphaproteobacteria |
| 99 | Mesorhizobium | Proteobacteria;Alphaproteobacteria |
| 100 | Mesorhizobium | Proteobacteria;Alphaproteobacteria |
| 101 | Methylobacterium | Proteobacteria;Alphaproteobacteria |
| 102 | Microbacteriaceae | Actinobacteria;Actinobacteria |
| 103 | Microbacterium | Actinobacteria;Actinobacteria |
| 104 | Microbacterium | Actinobacteria;Actinobacteria |
| 105 | Microbacterium | Actinobacteria;Actinobacteria |
| 106 | Microbacterium | Actinobacteria;Actinobacteria |
| 107 | Microbacterium | Actinobacteria;Actinobacteria |
| 108 | Microbacterium | Actinobacteria;Actinobacteria |
| 109 | Microbacterium | Actinobacteria;Actinobacteria |
| 110 | Microbacterium | Actinobacteria;Actinobacteria |
| 111 | Microbacterium | Actinobacteria;Actinobacteria |
| 112 | Microbacterium | Actinobacteria;Actinobacteria |
| 113 | Mucilaginibacter | Bacteroidetes;Bacteroidia |
| 114 | Mucilaginibacter | Bacteroidetes;Bacteroidia |
| 115 | Mucilaginibacter | Bacteroidetes;Bacteroidia |
| 116 | Mumia | Actinobacteria;Actinobacteria |
| 117 | Mycobacterium | Actinobacteria;Actinobacteria |
| 118 | Mycobacterium | Actinobacteria;Actinobacteria |
| 119 | Mycobacterium | Actinobacteria;Actinobacteria |
| 120 | Nocardioides | Actinobacteria;Actinobacteria |
| 121 | Nocardioides | Actinobacteria;Actinobacteria |
| 122 | Nocardioides | Actinobacteria;Actinobacteria |
| 123 | Nocardioides | Actinobacteria;Actinobacteria |
| 124 | Nocardioides | Actinobacteria;Actinobacteria |
| 125 | Nocardioides | Actinobacteria;Actinobacteria |
| 126 | Nocardioides | Actinobacteria;Actinobacteria |
| 127 | Nocardioides | Actinobacteria;Actinobacteria |
| 128 | Nocardioides | Actinobacteria;Actinobacteria |
| 129 | Nocardioides | Actinobacteria;Actinobacteria |
| 130 | Nocardioides | Actinobacteria;Actinobacteria |
| 131 | Nocardioides | Actinobacteria;Actinobacteria |
| 132 | Nonomuraea | Actinobacteria;Actinobacteria |
| 133 | Nonomuraea | Actinobacteria;Actinobacteria |
| 134 | Phenylobacterium | Proteobacteria;Alphaproteobacteria |
| 135 | Phycicoccus | Actinobacteria;Actinobacteria |
| 136 | Phycicoccus | Actinobacteria;Actinobacteria |
| 137 | Pseudomonas | Proteobacteria;Gammaproteobacteria |
| 138 | Pseudomonas | Proteobacteria;Gammaproteobacteria |
| 139 | Pseudomonas | Proteobacteria;Gammaproteobacteria |
| 140 | Pseudomonas | Proteobacteria;Gammaproteobacteria |
| 141 | Pseudomonas | Proteobacteria;Gammaproteobacteria |

| | | |
|---|---|---|
| 142 | Pseudoxanthomonas | Proteobacteria;Gammaproteobacteria |
| 143 | Pseudoxanthomonas | Proteobacteria;Gammaproteobacteria |
| 144 | Rhizobacter | Proteobacteria;Gammaproteobacteria |
| 145 | Rhizobacter | Proteobacteria;Gammaproteobacteria |
| 146 | Rhizobacter | Proteobacteria;Gammaproteobacteria |
| 147 | Rhizobiaceae | Proteobacteria;Alphaproteobacteria |
| 148 | Rhizobiaceae | Proteobacteria;Alphaproteobacteria |
| 149 | Rhizobiales | Proteobacteria;Alphaproteobacteria |
| 150 | Rhodococcus | Actinobacteria;Actinobacteria |
| 151 | Rhodococcus; | Actinobacteria;Actinobacteria |
| 152 | Roseateles | Proteobacteria;Gammaproteobacteria |
| 153 | Serratia | Proteobacteria;Gammaproteobacteria |
| 154 | Serratia | Proteobacteria;Gammaproteobacteria |
| 155 | Sphingomonadaceae;uncultured | Proteobacteria;Alphaproteobacteria |
| 156 | Sphingomonas | Proteobacteria;Alphaproteobacteria |
| 157 | Sphingomonas | Proteobacteria;Alphaproteobacteria |
| 158 | Sphingopyxis | Proteobacteria;Alphaproteobacteria |
| 159 | Sphingopyxis | Proteobacteria;Alphaproteobacteria |
| 160 | Stenotrophomonas | Proteobacteria;Gammaproteobacteria |
| 161 | Streptomyces | Actinobacteria;Actinobacteria |
| 162 | Streptomyces | Actinobacteria;Actinobacteria |
| 163 | Tardiphaga | Proteobacteria;Alphaproteobacteria |
| 164 | Variovorax | Proteobacteria;Gammaproteobacteria |
| 165 | Xanthobacteraceae | Proteobacteria;Alphaproteobacteria |
| 166 | Xanthobacteraceae;uncultured | Proteobacteria;Alphaproteobacteria |
| 167 | Unclassified; | Unclassified |
| 168 | Unclassified; | Unclassified |
| 169 | Unclassified; | Unclassified |

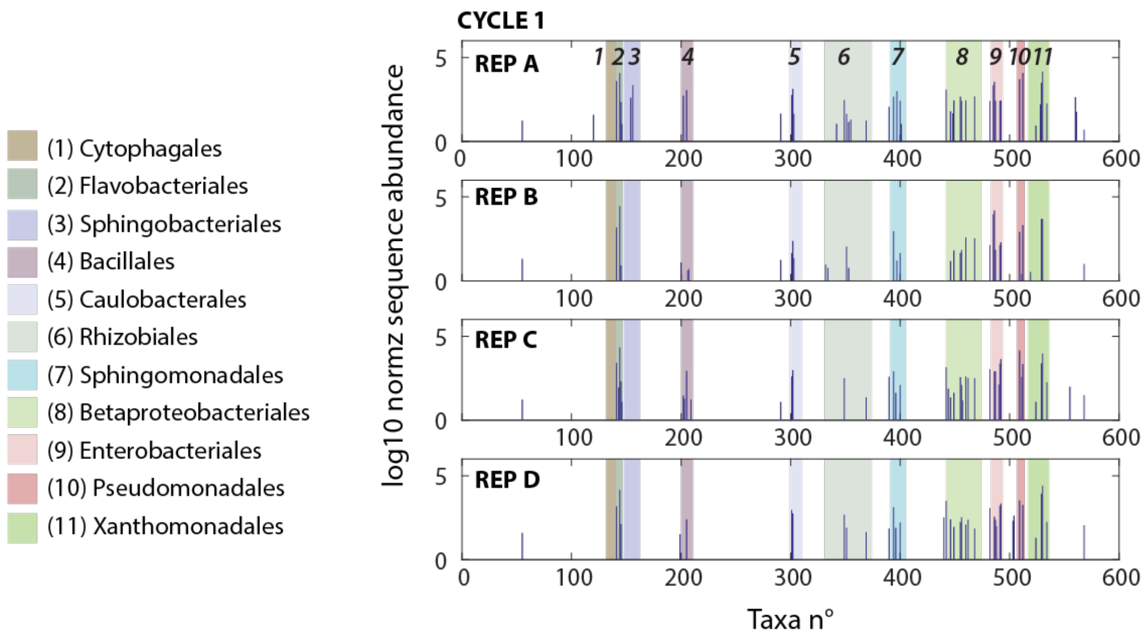


**Supplementary figure S1. Replicate taxa variation of NatComs after the first week growth cycle.** Plots show mean $log_{10}$-transformed total read-normalized ($5 \times 10^4$) taxa abundances in the soil inoculum, bars positioned according to taxa numbering from the OTU list (SILVA, above 99% similarity), with background color representing order affiliation (numbering and coloring, according to legend). Note how order attribution is maintained among replicates but exact OTU assignments within orders are varying among replicates.

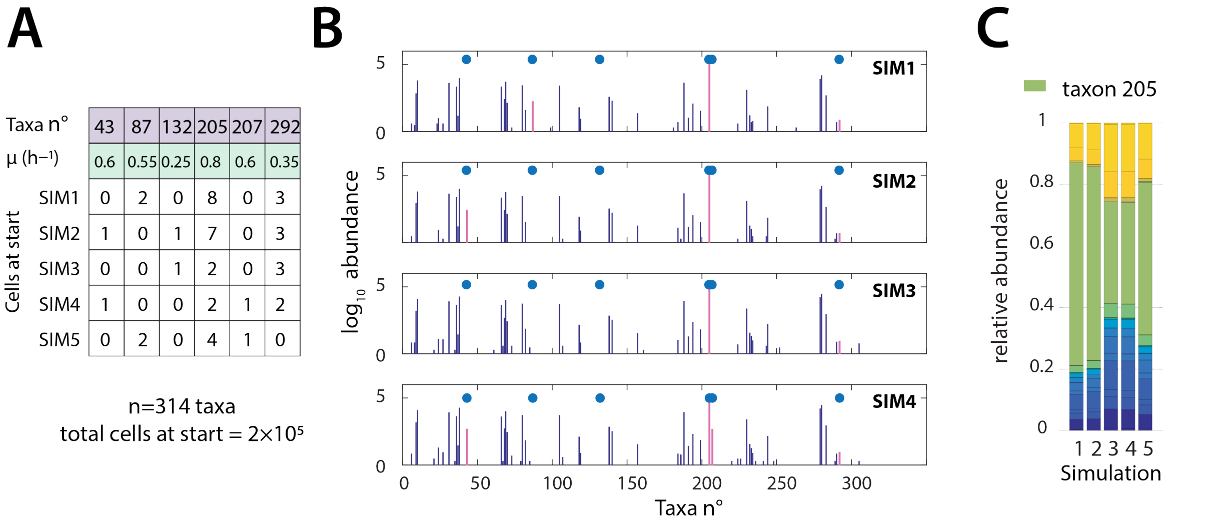


**Supplementary figure S2. Simulation of stochastic subsampling effects from a species-rich inoculum on the community composition after one week of growth. (A)** Communities are randomly subsampled to 200,000 cells from a soil community distribution with $n$ = 314 measured taxa and their relative abundances (*4*). Growth rates are assigned between 0.01–0.4 $h^{-1}$ skewed by the $log_{10}$-relative species abundance. Six low-abundance taxa with subsampled varying cell numbers at start in five simulations (SIM1-SIM5) between 0 and 10 are given high growth rates (0.25–0.8 $h^{-1}$). **(B)** Stationary phase taxa abundances in four replicate simulations (SIM 1–4) of multispecies growth according to the carbon-limited community growth model proposed in Ref. (*5*). Final community size after simulated growth attains ~$2\times10^8$ cells, from which 200,000 are randomly subsampled, summed per taxa number (as if for sequencing with $2\times10^5$ reads) and displayed (magenta bars and blue dots). Note the effects of variable starting cell numbers on the final abundances of the taxa numbers in A. **(C)** Stacked stationary phase relative abundances of the simulations of panel A and B, highlighting in green the taxon 205.

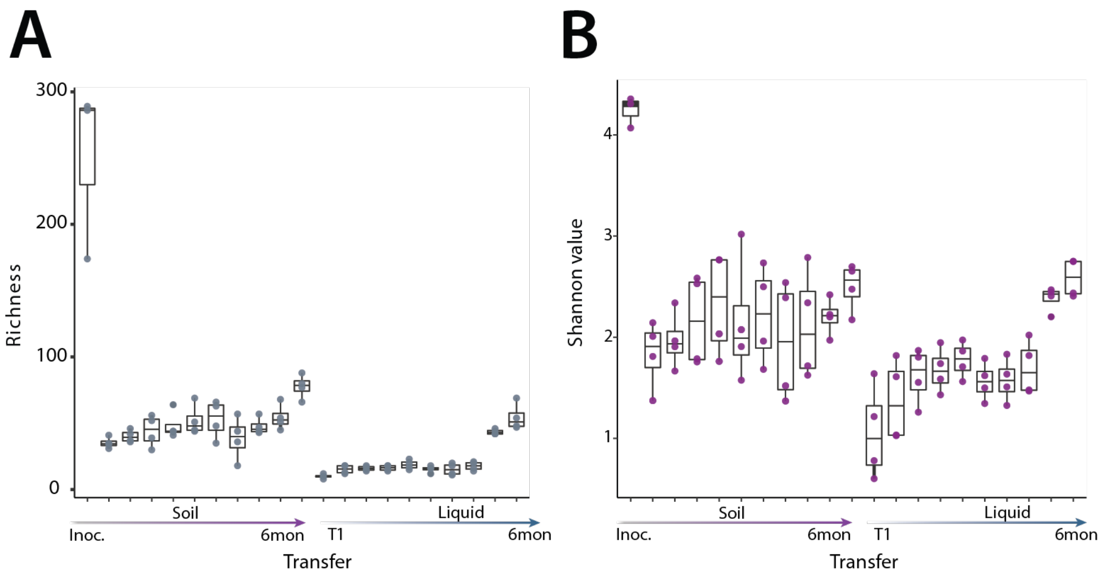


**Supplementary figure S3. Community diversity of NatComs in soil and liquid microcosms. (A)** Mean OTU richness (regular box plot, four replicates) of the NatCom inoculum and the successive communities after the one-week growth/dilution cycles in soil and liquid (T1–T8), 2 and 6 months (6mon) prolonged incubations (in that order of the Transfer arrow). Both soil and liquid supplemented with the same volume of soil extract as nutrient source. **(B)** As for (A) but the calculated Shannon index.

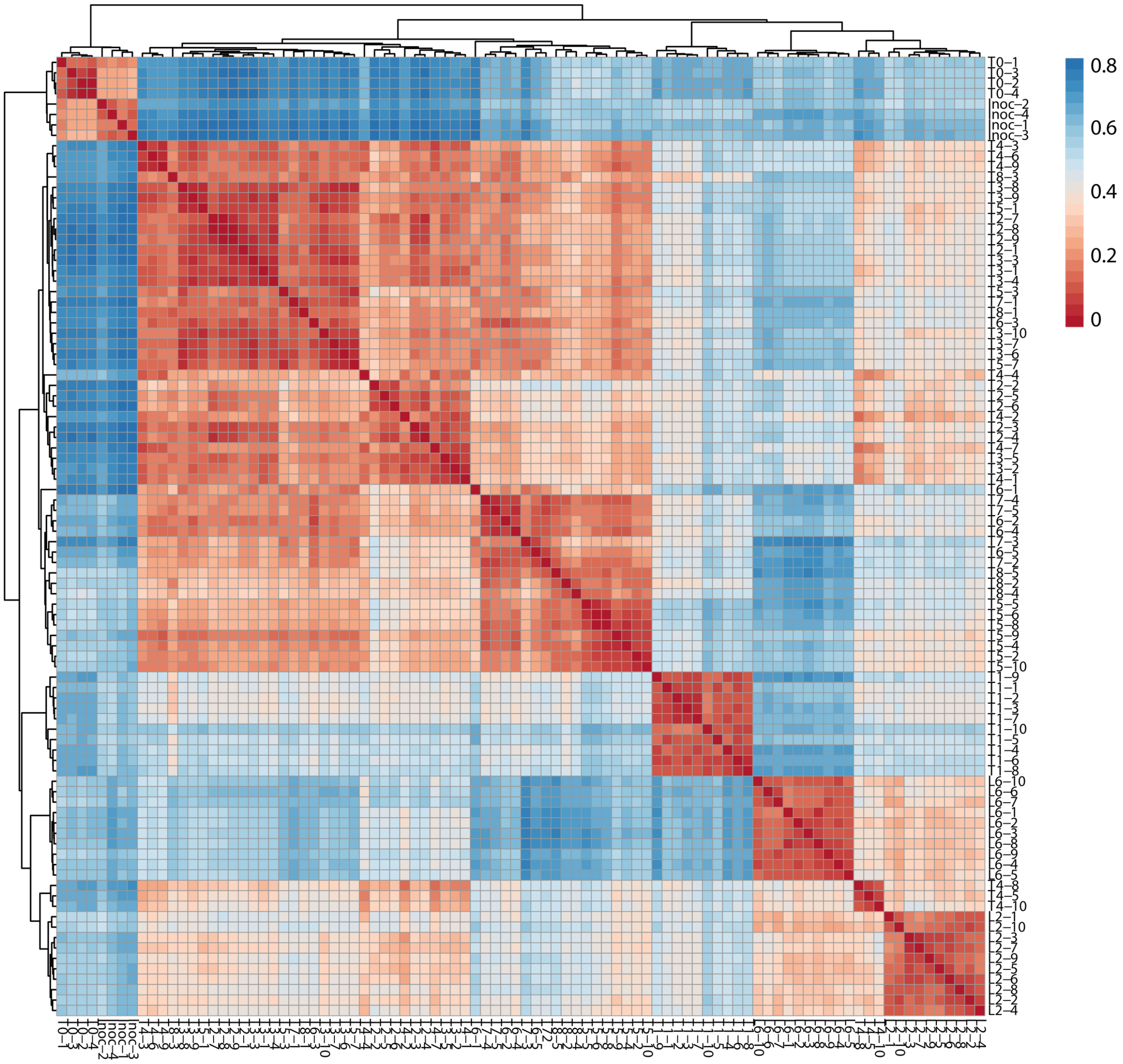


**Supplementary figure S4. Replicate community variability of the SynComs.** Plot shows pairwise sample comparisons, average-linkage clustered samples based on Bray-Curtis distances (color scale of the heatmap representation). Inoculum (Inoc 1–4), T0 samples (1–4, directly after addition), growth cycles (T1–T5, each in 10 replicates, T6-T8, each in 5 replicates), and prolonged incubations (L2, 2 mo; L6, 6 mo, each in 10 replicates) in small fonts on the right and bottom. Note the absence of replicate signature, but the presence of time sample signatures (e.g., L2 and L6).

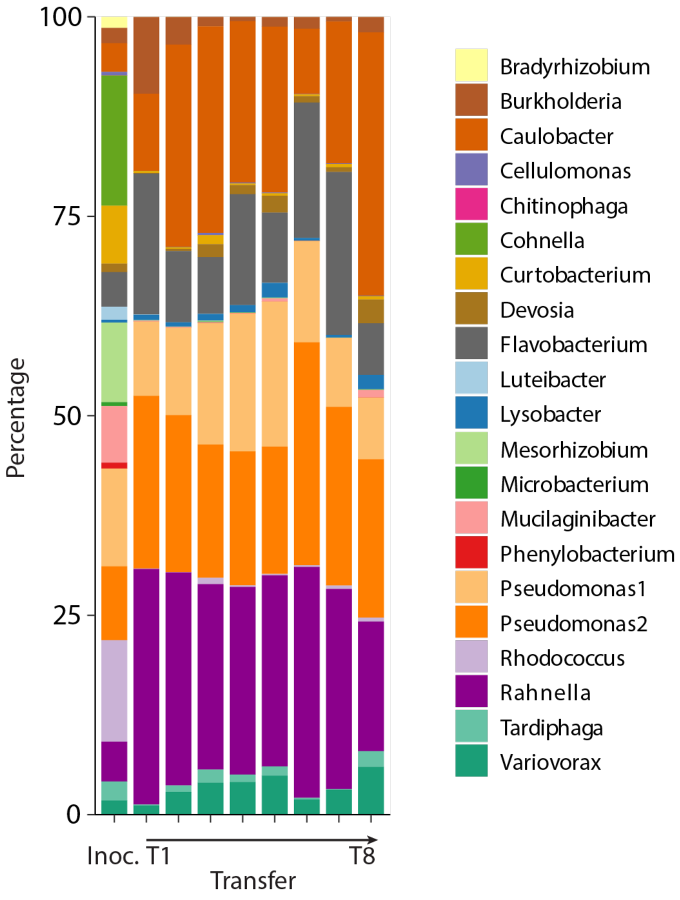


**Supplementary figure S5. SynCom diversity changes during growth/dilution cycles in liquid microcosms.** Plot shows stacked mean relative abundances per SynCom member from growth cycle T1 until T8 (*n* = 10 replicates for T1-T4, then 5 replicates for T5–T8), in comparison to the inoculum (Inoc.).

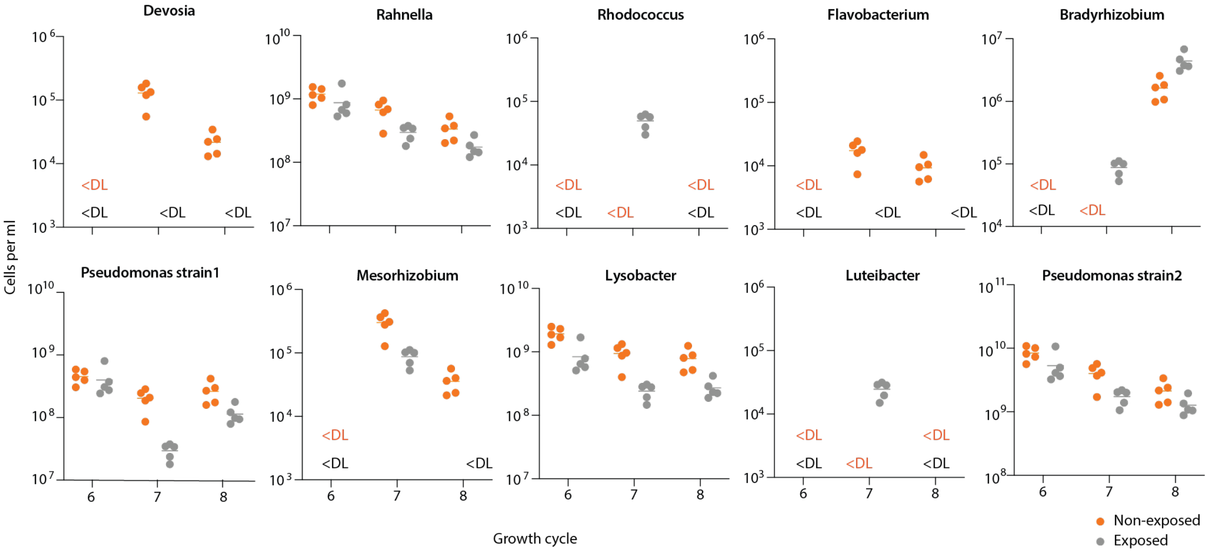


**Supplementary figure S6. Mean abundance changes of SynCom members upon toluene exposure.** Plots show mean absolute population sizes (calculated from individual relative sequence abundances and total community size by flow cytometry, *n* = 5 replicates) of SynCom members without (orange dots) and with toluene (grey dots) exposure. Plot shows strains not present in Fig. 6D. Strains with zero counts in both conditions are not shown.

### Supplementary References


1. C. A. Stedmon, S. Markager, R. Bro, Tracing dissolved organic matter in aquatic environments using a new approach to fluorescence spectroscopy. *Marine Chemistry* **82**, 239 (2003).10.1016/s0304-4203(03)00072-0
2. J. B. Fellman, E. Hood, R. G. M. Spencer, Fluorescence spectroscopy opens new windows into dissolved organic matter dynamics in freshwater ecosystems: A review. *Limnol Oceanogr* **55**, 2452 (2010).10.4319/lo.2010.55.6.2452
3. F. Behar, V. Beaumont, H. L. De B. Penteado, Rock-Eval 6 Technology: Performances and Developments. *Oil & Gas Science and Technology* **56**, 111 (2006).10.2516/ogst:2001013
4. M. Dubey, N. Hadadi, S. Pelet, N. Carraro, D. R. Johnson, J. R. van der Meer, Environmental connectivity controls diversity in soil microbial communities. *Commun Biol* **4**, 492 (2021).10.1038/s42003-021-02023-2
5. N. Hadadi, J. R. van der Meer. (Zenodo, 2021). http://doi.org/10.5281/zenodo.4568347
